# DNA Replication-transcription conflicts do not significantly contribute to spontaneous mutations due to replication errors in *Escherichia coli*.

**DOI:** 10.1101/2021.08.23.457454

**Authors:** Patricia L. Foster, Brittany A. Niccum, Heewook Lee

**Affiliations:** Department of Biology, Indiana University, Bloomington, IN, 47405, USA; Luddy School of Informatics, Computing and Engineering, Indiana University, Bloomington, IN, 47405, USA

**Keywords:** Replication-transcription conflicts, mutation accumulation, mismatch repair, base-pair substitutions, indels

## Abstract

Encounters between DNA replication and transcription can cause genomic disruption, particularly when the two meet head-on. Whether these conflicts produce point mutations is debated. This paper presents detailed analyses of a large collection of mutations generated during mutation accumulation experiments with mismatch-repair (MMR) defective *Escherichia coli*. With MMR absent, mutations are primarily due to DNA replication errors. Overall, there were no differences in the frequencies of base-pair substitutions or small indels (insertion and deletions ≤ 4 bp) in the coding sequences or promoters of genes oriented codirectionally versus head-on to replication. Among a subset of highly expressed genes there was a 2- to 3-fold bias for indels in genes oriented head-on to replication, but this difference was almost entirely due to the asymmetrical genomic locations of tRNA genes containing mononucleotide runs, which are hotspots for indels. No additional orientation bias in mutation frequencies occurred when MMR^−^ strains were also defective for transcription-coupled repair (TCR). However, in contrast to other reports, loss of TCR slightly increased the overall mutation rate, meaning that TCR is antimutagenic. There was no orientation bias in mutation frequencies among the stress-response genes that are regulated by RpoS or induced by DNA damage. Thus, biases in the locations of mutational targets can account for most, if not all, apparent biases in mutation frequencies between genes oriented head-on versus co-directional to replication. In addition, the data revealed a strong correlation of the frequency of base-pair substitutions with gene length, but no correlation with gene expression levels.

**IMPORTANCE:** Because DNA replication and transcription occur on the same DNA template, encounters between the two machines occur frequently. When these encounters are head-to-head, genomic disruption can occur. But, whether replication-transcription conflicts contribute to spontaneous mutations is debated. Analyzing in detail a large collection of mutations generated with mismatch repair defective *Escherichia coli* strains, we find that across the genome there are no significant differences in mutation frequencies between genes oriented co-directionally versus head-on to replication. Among a subset of highly expressed genes there was a 2- to 3-fold bias for small insertions and deletions in head-on oriented genes, but this difference was almost entirely due to the asymmetrical locations of tRNA genes containing mononucleotide runs, which are hotspots for these mutations. Thus, biases in the positions of mutational target sequences can account for most, if not all, apparent biases in mutation frequencies between genes oriented head-on and co-directionally to replication.

## INTRODUCTION

The fact that DNA replication and transcription occur simultaneously and on the same template sets up inevitable molecular conflicts. In *Escherichia coli* replication proceeds at an average of 650 bps·per second (1) whereas transcription, at its fastest, proceeds at about a tenth of this speed (2). Also, since there are thousands of RNA polymerase molecules active at a time (3), collisions between the replication fork and the transcription complex must be frequent. The effects of these collisions on genomic integrity have been a subject of active research for years. In particular, the mutagenic and evolutionary consequences of collisions that occur when replication and transcription meet head-on versus codirectionally is the subject of recent debate in the literature (4, 5).

Head-on (HO) collisions occur when the transcribed DNA strand is the template for lagging-strand replication, whereas codirectional (CD) collisions occur when the transcribed DNA strand is the template for leading-strand replication. HO, but not CD collisions, slow replication in bacteria (6-8) and yeast (9). In addition, experiments using experimentally inverted highly-expressed genes show that HO collisions can arrest replication forks and produce DNA double-strand breaks, requiring recombination functions to restore replication (10). The potential of HO replication-transcription conflicts to produce genomic disruption is a powerful selective force, one indication of which is that ribosomal operons, in which genes are highly expressed, are in the CD orientation in almost all bacteria (11). Many bacteria also have a large majority of all their genes in the CD orientation, although *E. coli* is an exception to this tendency with only 55% of its genes so oriented (12).

While clearly a threat to overall genomic maintenance and stability, it is not as clear that replication-transcription conflicts normally contribute to spontaneous point-mutation rates. Two types of evidence suggest that they do. First, comparative genomics of diverged strains of *B. subtilis* showed HO-oriented genes have a 42% higher rate of non-synonymous (amino-acid changing) substitutions than do CD-oriented genes, although the rate of synonymous substitutions is nearly the same (13). Second, when a gene is experimentally inverted, its mutation frequency can be greater in the HO orientation than in the CD orientation (8, 13-15). However, these observations have alternative explanations (5), which will be further considered in the discussion section.

Several reports have emphasized that the mutations induced by replication-transcription conflicts have signatures [reviewed in (5)]. In both *E. coli* (16) and *B. subtilis* (15), inverting a gene revealed orientation-specific hotspots for both BPSs and large duplications and deletions. Interestingly, in both studies the HO conflict promoted a specific base-pair substitution (BPS) in the -10 element of the gene’s promoter (15, 16).

In a previous publication (17) we reported that there was no difference in overall BPS rates of HO versus CD oriented genes in *E. coli*. This conclusion was based on mutation accumulation (MA) experiments followed by whole-genome sequencing (WGS) that yielded 233 BPS mutations in a wild-type strain and 1625 BPS mutations in a mismatch repair (MMR) defective strain (18). Since our original publication we have conducted nine additional MA-WGS experiments with MMR defective strains, yielding a total of 30,061 BPSs and 5324 small insertions or deletions (indels) (19). MMR corrects mismatches in newly replicated DNA [reviewed in (20)], thus these mutations can be considered to be primarily errors made during normal replication. For this report we used this extensive database to investigate in greater detail whether replication-transcription conflicts induce the types of mutations detected in other experiments, or, possibly, other characteristic mutations. Our conclusion is that in *E. coli* growing and replicating in rich medium and free of exogenous stress, replication-transcription conflicts are not a major source of spontaneous point mutations.

## RESULTS

### Distributions of mutation frequencies per gene and correlations to other parameters

Combining the results of ten MA-WGS experiments with MMR^—^ strains gave a database of 30,061 BPSs and 5324 indels (≤ 4 bp) across the *E. coli* genome (19). Of these, 27,164 BPSs and 3841 indels were in gene coding sequences (CDSs) (see Table S1 and Table S2 for the primary data discussed below). Before applying standard statistical methods to these data, we analyzed the distributions of the mutation frequencies. If the numbers of mutations/CDS were random, the distribution would be Poisson, but the BPSs/CDS numbers fell into a normal (Gaussian) distribution (Fig. 1A). As was found for *B. subtilis* (21), there is a strong linear correlation between the number of BPSs in a CDS and the length of the CDS in nucleotides (Nts) (Fig. 1B & 1C and Table 1). Fig. 1D shows that CDS length is normally distributed except for the ribosomal and tRNA genes (which will be further discussed below). These two factors explain the normal distribution of BPSs/CDS.

**Figure 1.**
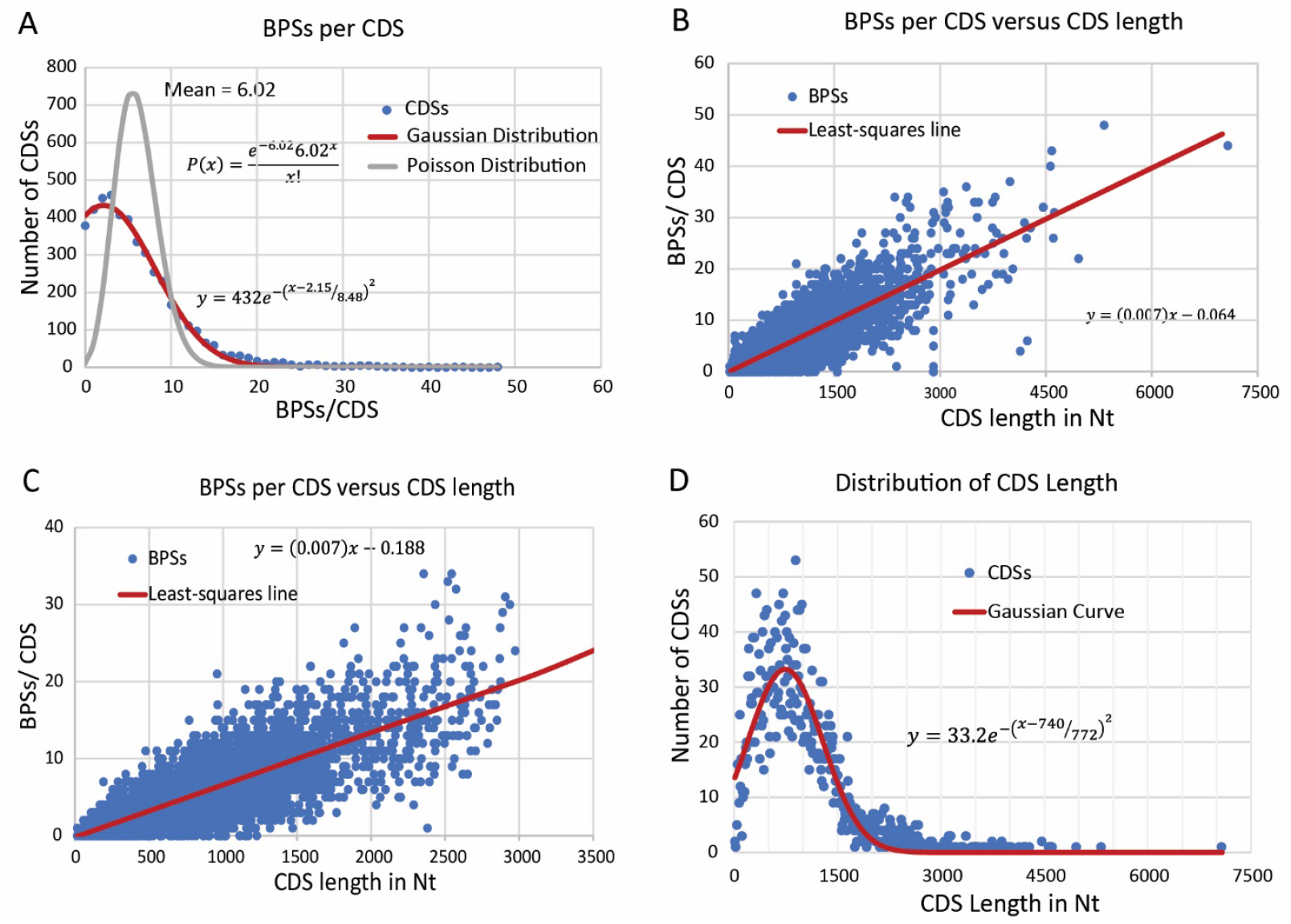
The distributions of the numbers of base-pair substitutions (BPSs) and the lengths of gene coding sequences (CDSs). (A) The distribution of the BPSs per CDS. The numbers of gene CDSs (Y-axis) containing the indicated numbers of BPSs (X axis) is shown (blue circles). A Poisson distribution for the mean of 6.02 (gray line) clearly does not describe the data but a Gaussian curve (red line) does. The Gaussian equation shown gives a fit with the coefficient of determination, R^2^, equal to 0.995. (B) The relationship of BPSs per CDS and length of the CDS in Nts. The number of BPSs/CDS (blue circles) is plotted against the length of the CDS in nucleotides (Nts). The equation shows the least-squares fit (red line), which gives a correlation coefficient, R, of 0.81 (P<0.0003). (C) The same as (B), but to show that the correlation does not depend on outliers, CDSs longer that 3000 Nt and the ribosomal and tRNA genes have been eliminated (see text for explanation for eliminating these genes). The equation shows the least-squares fit (red line), which gives a correlation coefficient, R, of 0.78 (P<0.0003). (D) The distribution of CDS lengths. The number of CDSs (Y-axis) of the indicated length (X axis) is shown (blue circles). For this analysis, the CDS lengths were binned into 10 Nt long bins. A Gaussian curve (red line) was fit to the data disregarding ribosomal and tRNA genes (see text for explanation for eliminating these genes). The Gaussian equation shown gives a fit with the coefficient of determination, R^2^, equal to 0.89.

**Table 1.**
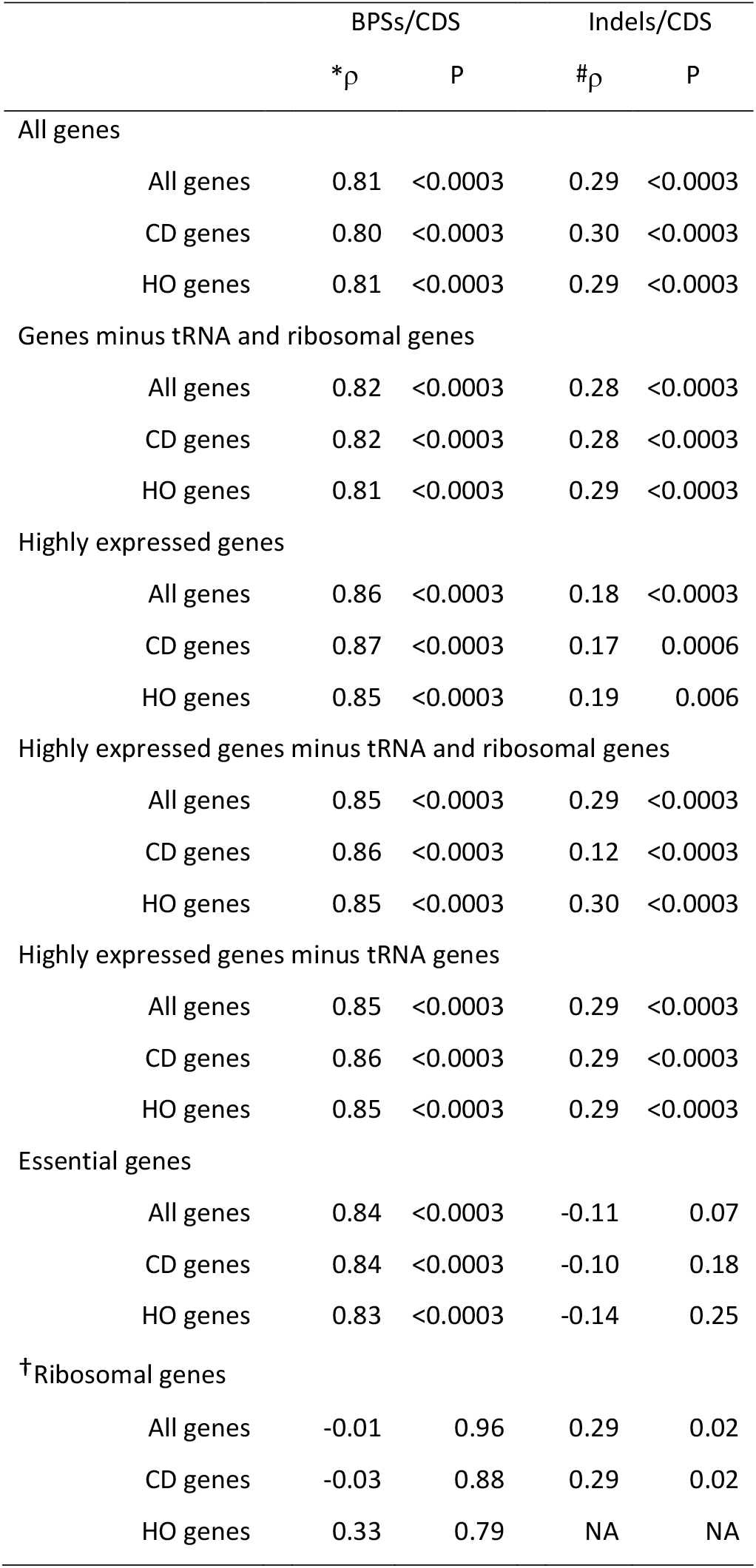
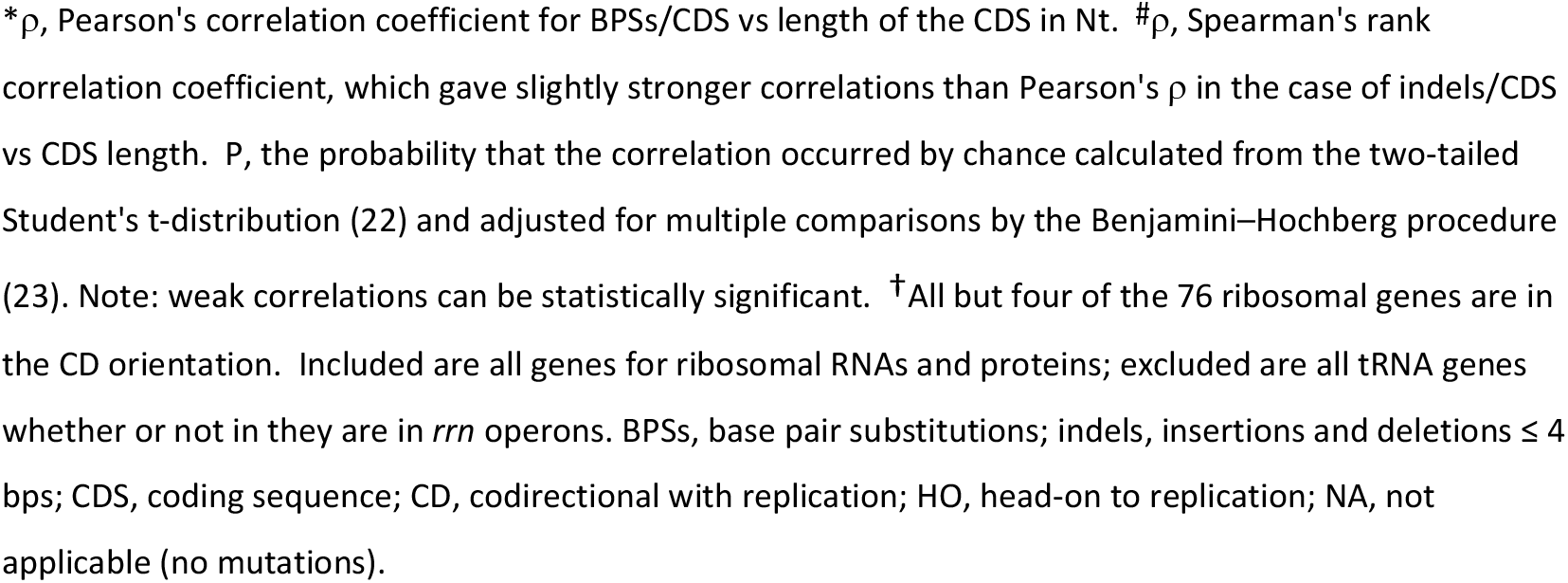
Correlations of mutations per CDS with CDS length

|  |  | BPSs/CDS |  | Indels/CDS |  |
| --- | --- | --- | --- | --- | --- |
| | | *$\rho$ | P | #$\rho$ | P |
| All genes |  |  |  |  |  |
|  | All genes | 0.81 | <0.0003 | 0.29 | <0.0003 |
|  | CD genes | 0.80 | <0.0003 | 0.30 | <0.0003 |
|  | HO genes | 0.81 | <0.0003 | 0.29 | <0.0003 |
| Genes minus tRNA and ribosomal genes |  |  |  |  |  |
|  | All genes | 0.82 | <0.0003 | 0.28 | <0.0003 |
|  | CD genes | 0.82 | <0.0003 | 0.28 | <0.0003 |
|  | HO genes | 0.81 | <0.0003 | 0.29 | <0.0003 |
| Highly expressed genes |  |  |  |  |  |
|  | All genes | 0.86 | <0.0003 | 0.18 | <0.0003 |
|  | CD genes | 0.87 | <0.0003 | 0.17 | 0.0006 |
|  | HO genes | 0.85 | <0.0003 | 0.19 | 0.006 |
| Highly expressed genes minus tRNA and ribosomal genes |  |  |  |  |  |
|  | All genes | 0.85 | <0.0003 | 0.29 | <0.0003 |
|  | CD genes | 0.86 | <0.0003 | 0.12 | <0.0003 |
|  | HO genes | 0.85 | <0.0003 | 0.30 | <0.0003 |
| Highly expressed genes minus tRNA genes |  |  |  |  |  |
|  | All genes | 0.85 | <0.0003 | 0.29 | <0.0003 |
|  | CD genes | 0.86 | <0.0003 | 0.29 | <0.0003 |
|  | HO genes | 0.85 | <0.0003 | 0.29 | <0.0003 |
| Essential genes |  |  |  |  |  |
|  | All genes | 0.84 | <0.0003 | -0.11 | 0.07 |
|  | CD genes | 0.84 | <0.0003 | -0.10 | 0.18 |
|  | HO genes | 0.83 | <0.0003 | -0.14 | 0.25 |
| <sup>†</sup>Ribosomal genes |  |  |  |  |  |
|  | All genes | -0.01 | 0.96 | 0.29 | 0.02 |
|  | CD genes | -0.03 | 0.88 | 0.29 | 0.02 |
|  | HO genes | 0.33 | 0.79 | NA | NA |

\* $\rho$ , Pearson's correlation coefficient for BPSs/CDS vs length of the CDS in Nt. # $\rho$ , Spearman's rank correlation coefficient, which gave slightly stronger correlations than Pearson's $\rho$ in the case of indels/CDS vs CDS length. P, the probability that the correlation occurred by chance calculated from the two-tailed Student's t-distribution (22) and adjusted for multiple comparisons by the Benjamini–Hochberg procedure (23). Note: weak correlations can be statistically significant. †All but four of the 76 ribosomal genes are in the CD orientation. Included are all genes for ribosomal RNAs and proteins; excluded are all tRNA genes whether or not in they are in *rrn* operons. BPSs, base pair substitutions; indels, insertions and deletions $\leq 4$ bps; CDS, coding sequence; CD, codirectional with replication; HO, head-on to replication; NA, not applicable (no mutations).

In contrast to the BPSs, the numbers of indels/CDS fell into neither a Poisson nor a normal distribution. Because indel mutation rate is dependent on the length of homopolymeric runs (18, 24), and the occurrence of such runs in genes appears to be idiosyncratic, the distribution of indels/CDS is also idiosyncratic.

We tested for correlations between mutations/CDS and mutations/CDS/Nt and other parameters. There were no significant differences between the frequencies of mutations in CDSs located in the right or left replichores, or between CDSs in the forward or reverse orientation relative to the reference strand. The sole exception was a small (∼20%) greater value of BPSs/CDS/Nt among highly expressed genes (minus ribosomal and tRNA genes) in the left replichore versus those in right replichore. There was also no significant correlation between mutation frequencies and distance of a CDS from the origin, but the symmetrical wave-pattern of mutation rates across the genome, previously reported (25, 26), would not have been detected as a monotonic correlation. Thus, the correlation between BPSs/CDS and length of the CDS remained the only significant correlation overall. (Data not shown but available in the IUScholarWorks Repository http://hdl.handle.net/2022/25294.)

It is generally agreed that transcription is mutagenic and, consequently, highly expressed genes have greater mutation rates than poorly expressed genes [reviewed in (27)]. However, the opposite has also been reported (28-30). We used our previously published RNA-seq values from lag, log, and stationary phase cells (26) to determine if mutation frequencies were correlated to gene expression levels. Surprisingly, we found no overall correlation (Table 2). An exception was the subset of genes encoding the ribosomal RNAs and proteins, which showed weak correlations between BPSs/CDS and gene expression levels.

**Table 2.**
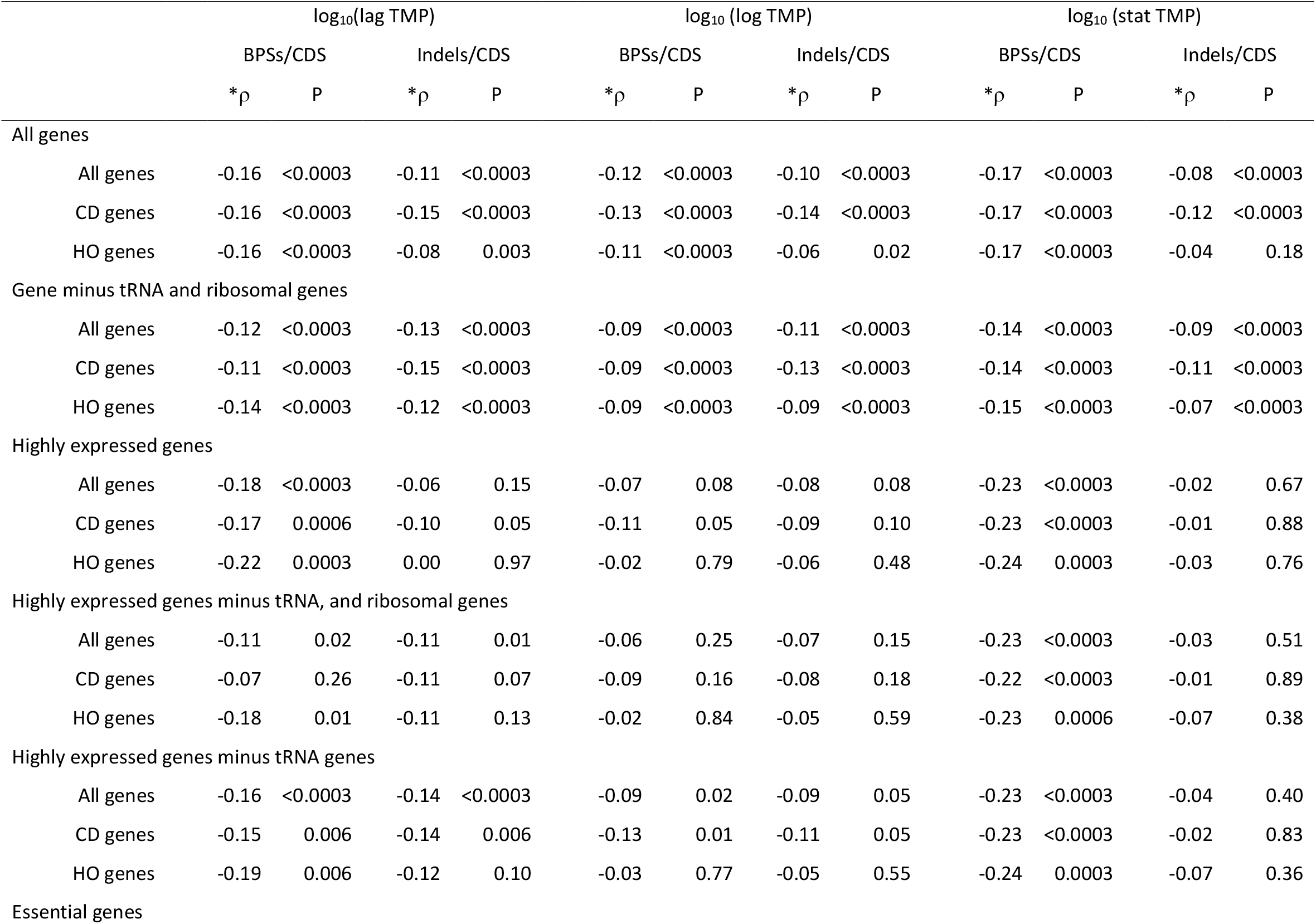
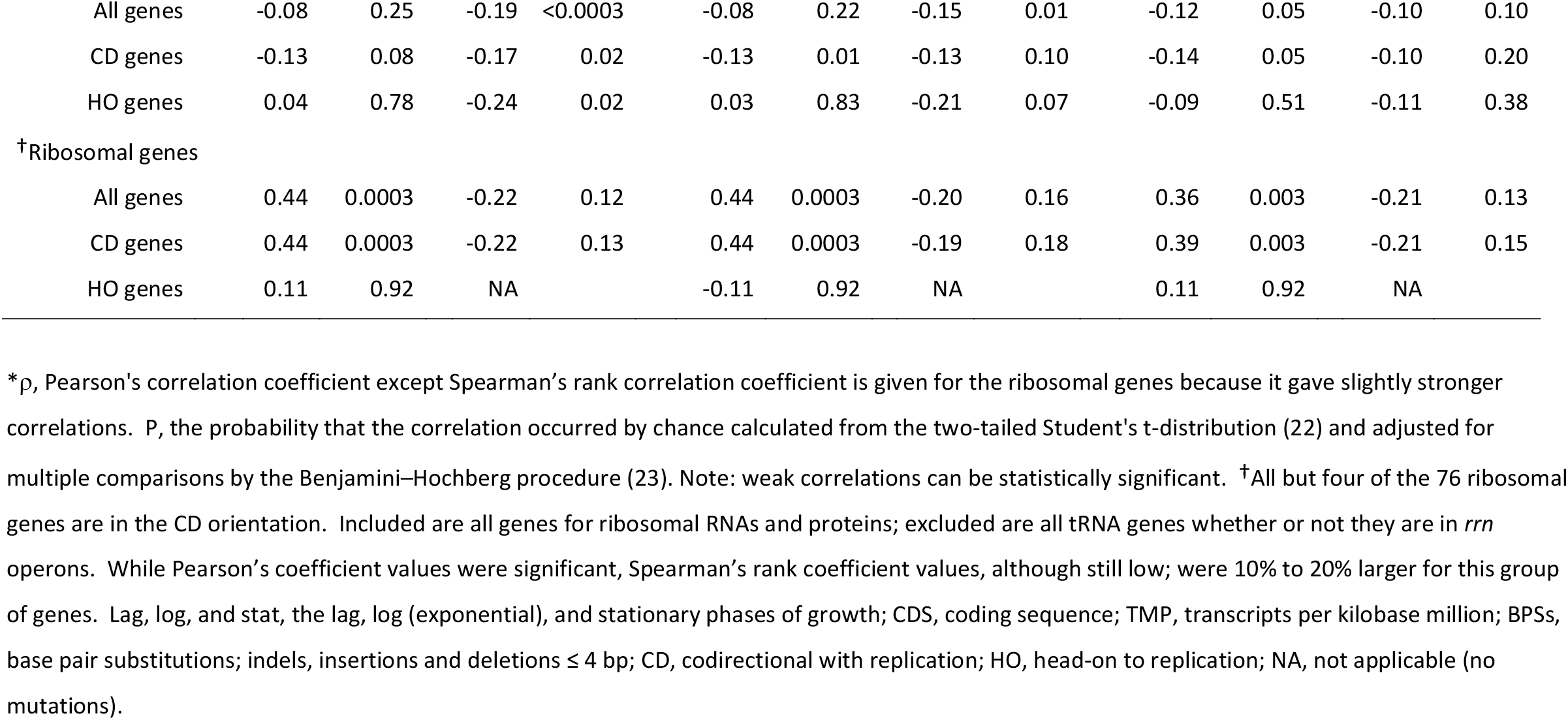
Correlations of the frequencies of mutations per CDS with gene expression levels

| | | $\log_{10}(\text{lag TMP})$ | | | | $\log_{10}(\text{log TMP})$ | | | | $\log_{10}(\text{stat TMP})$ | | | |
| --- | --- | --- | --- | --- | --- | --- | --- | --- | --- | --- | --- | --- | --- |
|  |  | BPSs/CDS |  | Indels/CDS |  | BPSs/CDS |  | Indels/CDS |  | BPSs/CDS |  | Indels/CDS |  |
| | | *$\rho$ | P | *$\rho$ | P | *$\rho$ | P | *$\rho$ | P | *$\rho$ | P | *$\rho$ | P |
| All genes |  |  |  |  |  |  |  |  |  |  |  |  |  |
|  | All genes | -0.16 | <0.0003 | -0.11 | <0.0003 | -0.12 | <0.0003 | -0.10 | <0.0003 | -0.17 | <0.0003 | -0.08 | <0.0003 |
|  | CD genes | -0.16 | <0.0003 | -0.15 | <0.0003 | -0.13 | <0.0003 | -0.14 | <0.0003 | -0.17 | <0.0003 | -0.12 | <0.0003 |
|  | HO genes | -0.16 | <0.0003 | -0.08 | 0.003 | -0.11 | <0.0003 | -0.06 | 0.02 | -0.17 | <0.0003 | -0.04 | 0.18 |
| Gene minus tRNA and ribosomal genes |  |  |  |  |  |  |  |  |  |  |  |  |  |
|  | All genes | -0.12 | <0.0003 | -0.13 | <0.0003 | -0.09 | <0.0003 | -0.11 | <0.0003 | -0.14 | <0.0003 | -0.09 | <0.0003 |
|  | CD genes | -0.11 | <0.0003 | -0.15 | <0.0003 | -0.09 | <0.0003 | -0.13 | <0.0003 | -0.14 | <0.0003 | -0.11 | <0.0003 |
|  | HO genes | -0.14 | <0.0003 | -0.12 | <0.0003 | -0.09 | <0.0003 | -0.09 | <0.0003 | -0.15 | <0.0003 | -0.07 | <0.0003 |
| Highly expressed genes |  |  |  |  |  |  |  |  |  |  |  |  |  |
|  | All genes | -0.18 | <0.0003 | -0.06 | 0.15 | -0.07 | 0.08 | -0.08 | 0.08 | -0.23 | <0.0003 | -0.02 | 0.67 |
|  | CD genes | -0.17 | 0.0006 | -0.10 | 0.05 | -0.11 | 0.05 | -0.09 | 0.10 | -0.23 | <0.0003 | -0.01 | 0.88 |
|  | HO genes | -0.22 | 0.0003 | 0.00 | 0.97 | -0.02 | 0.79 | -0.06 | 0.48 | -0.24 | 0.0003 | -0.03 | 0.76 |
| Highly expressed genes minus tRNA, and ribosomal genes |  |  |  |  |  |  |  |  |  |  |  |  |  |
|  | All genes | -0.11 | 0.02 | -0.11 | 0.01 | -0.06 | 0.25 | -0.07 | 0.15 | -0.23 | <0.0003 | -0.03 | 0.51 |
|  | CD genes | -0.07 | 0.26 | -0.11 | 0.07 | -0.09 | 0.16 | -0.08 | 0.18 | -0.22 | <0.0003 | -0.01 | 0.89 |
|  | HO genes | -0.18 | 0.01 | -0.11 | 0.13 | -0.02 | 0.84 | -0.05 | 0.59 | -0.23 | 0.0006 | -0.07 | 0.38 |
| Highly expressed genes minus tRNA genes |  |  |  |  |  |  |  |  |  |  |  |  |  |
|  | All genes | -0.16 | <0.0003 | -0.14 | <0.0003 | -0.09 | 0.02 | -0.09 | 0.05 | -0.23 | <0.0003 | -0.04 | 0.40 |
|  | CD genes | -0.15 | 0.006 | -0.14 | 0.006 | -0.13 | 0.01 | -0.11 | 0.05 | -0.23 | <0.0003 | -0.02 | 0.83 |
|  | HO genes | -0.19 | 0.006 | -0.12 | 0.10 | -0.03 | 0.77 | -0.05 | 0.55 | -0.24 | 0.0003 | -0.07 | 0.36 |
| Essential genes |  |  |  |  |  |  |  |  |  |  |  |  |  |

|  |  |  |  |  |  |  |  |  |  |  |  |  |
| --- | --- | --- | --- | --- | --- | --- | --- | --- | --- | --- | --- | --- |
| All genes | -0.08 | 0.25 | -0.19 | <0.0003 | -0.08 | 0.22 | -0.15 | 0.01 | -0.12 | 0.05 | -0.10 | 0.10 |
| CD genes | -0.13 | 0.08 | -0.17 | 0.02 | -0.13 | 0.01 | -0.13 | 0.10 | -0.14 | 0.05 | -0.10 | 0.20 |
| HO genes | 0.04 | 0.78 | -0.24 | 0.02 | 0.03 | 0.83 | -0.21 | 0.07 | -0.09 | 0.51 | -0.11 | 0.38 |
| <sup>†</sup>Ribosomal genes |  |  |  |  |  |  |  |  |  |  |  |  |
| All genes | 0.44 | 0.0003 | -0.22 | 0.12 | 0.44 | 0.0003 | -0.20 | 0.16 | 0.36 | 0.003 | -0.21 | 0.13 |
| CD genes | 0.44 | 0.0003 | -0.22 | 0.13 | 0.44 | 0.0003 | -0.19 | 0.18 | 0.39 | 0.003 | -0.21 | 0.15 |
| HO genes | 0.11 | 0.92 | NA |  | -0.11 | 0.92 | NA |  | 0.11 | 0.92 | NA |  |

\* $\rho$ , Pearson's correlation coefficient except Spearman's rank correlation coefficient is given for the ribosomal genes because it gave slightly stronger correlations. P, the probability that the correlation occurred by chance calculated from the two-tailed Student's t-distribution (22) and adjusted for multiple comparisons by the Benjamini–Hochberg procedure (23). Note: weak correlations can be statistically significant. <sup>†</sup>All but four of the 76 ribosomal genes are in the CD orientation. Included are all genes for ribosomal RNAs and proteins; excluded are all tRNA genes whether or not they are in *rrn* operons. While Pearson's coefficient values were significant, Spearman's rank coefficient values, although still low; were 10% to 20% larger for this group of genes. Lag, log, and stat, the lag, log (exponential), and stationary phases of growth; CDS, coding sequence; TMP, transcripts per kilobase million; BPSs, base pair substitutions; indels, insertions and deletions $\leq 4$ bp; CD, codirectional with replication; HO, head-on to replication; NA, not applicable (no mutations).

### Overall, neither BPS nor indel frequencies are biased by gene orientation relative to DNA replication

Relative to the reference DNA strand, genes oriented in the forward direction on the right replichore and the reverse direction on the left replichore are transcribed codirectionally (CD) with replication, whereas genes oppositely oriented are transcribed head-on (HO) to replication. For the purposes of analysis, we set the division between the replichores half-way around the genome from the origin close to the *terC* site, the most frequent site of replication fork termination (31). This resulted in 2,467 CD-oriented genes and 2,044 HO-oriented genes. Table 3 shows there is no statistically significant difference in frequencies of BPSs or indels between CDS in the CD versus the HO orientation whether or not the frequencies are corrected for CDS length.

**Table 3.** Comparisons of the frequencies of mutations in CDSs oriented CD versus HO to replication

|  | BPSs/CDS |  | BPSs/CDS/Nt X 10<sup>3</sup> |  | Indels/CDS |  | Indels/CDS/Nt X 10<sup>3</sup> |  |
| --- | --- | --- | --- | --- | --- | --- | --- | --- |
|  | Mean | SD | Mean | SD | Mean | SD | Mean | SD |
| All genes |  |  |  |  |  |  |  |  |
| All genes | 6.02 | 5.33 | 6.54 | 4.46 | 0.85 | 2.32 | 1.30 | 9.32 |
| CD genes | 6.04 | 5.37 | 6.54 | 4.20 | 0.86 | 2.07 | 1.14 | 6.44 |
| HO genes | 6.01 | 5.28 | 6.55 | 4.76 | 0.84 | 2.60 | 1.49 | 11.9 |
| *ΔHO | 0% |  | 0% |  | -2% |  | 30% |  |
| #p | 0.90 |  | 0.97 |  | 0.88 |  | 0.36 |  |
| Genes minus tRNA and ribosomal genes |  |  |  |  |  |  |  |  |
| All genes | 6.19 | 5.34 | 6.48 | 3.89 | 0.85 | 2.24 | 1.03 | 3.51 |
| CD genes | 6.27 | 5.39 | 6.52 | 3.76 | 0.89 | 2.06 | 1.08 | 3.74 |
| HO genes | 6.10 | 5.28 | 6.44 | 4.04 | 0.81 | 2.44 | 0.97 | 3.22 |
| *ΔHO | -3% |  | -1% |  | -9% |  | -10% |  |
| #p | 0.40 |  | 0.64 |  | 0.38 |  | 0.43 |  |
| Highly expressed genes |  |  |  |  |  |  |  |  |
| All genes | 4.79 | 4.88 | 7.19 | 6.42 | 0.54 | 2.19 | 2.46 | 21.2 |
| CD genes | 4.81 | 4.88 | 7.08 | 5.54 | 0.40 | 1.62 | 1.19 | 12.3 |
| HO genes | 4.76 | 4.90 | 7.39 | 7.69 | 0.78 | 2.89 | 4.60 | 30.8 |
| *ΔHO | -1% |  | 4% |  | 92% |  | 287% |  |
| #p | 0.92 |  | 0.69 |  | 0.10 |  | 0.13 |  |
| Highly expressed genes minus tRNA, and ribosomal genes |  |  |  |  |  |  |  |  |
| All genes | 5.31 | 5.05 | 6.84 | 4.51 | 0.46 | 1.35 | 0.86 | 3.90 |
| CD genes | 5.44 | 5.09 | 6.94 | 4.66 | 0.43 | 1.37 | 0.77 | 3.09 |
| HO genes | 5.10 | 4.98 | 6.68 | 4.26 | 0.51 | 1.31 | 1.01 | 4.87 |
| *ΔHO | -6% |  | -4% |  | 17% |  | 31% |  |
| #p | 0.52 |  | 0.60 |  | 0.62 |  | 0.61 |  |
| Highly expressed genes minus tRNA genes |  |  |  |  |  |  |  |  |
| All genes | 5.10 | 4.93 | 6.83 | 4.56 | 0.43 | 1.30 | 0.80 | 3.75 |
| CD genes | 5.14 | 4.92 | 6.94 | 4.73 | 0.38 | 1.29 | 0.69 | 2.93 |
| HO genes | 5.02 | 4.96 | 6.65 | 4.27 | 0.50 | 1.30 | 0.99 | 4.83 |
| *ΔHO | -2% |  | -4% |  | 29% |  | 43% |  |
| #P | 0.84 |  | 0.54 |  | 0.39 |  | 0.50 |  |
| Essential genes |  |  |  |  |  |  |  |  |
| All genes | 6.23 | 5.55 | 6.34 | 3.56 | 0.09 | 0.50 | 0.23 | 1.50 |
| CD genes | 6.37 | 5.60 | 6.54 | 3.73 | 0.06 | 0.33 | 0.21 | 1.56 |
| HO genes | 5.92 | 5.44 | 5.84 | 3.10 | 0.15 | 0.75 | 0.27 | 1.37 |
| *ΔHO | -7% |  | -11% |  | 154% |  | 29% |  |
| #P | 0.61 |  | 0.13 |  | 0.35 |  | 0.82 |  |
| †Ribosomal genes |  |  |  |  |  |  |  |  |
| All genes | 2.18 | 2.23 | 5.51 | 6.76 | 0.17 | 0.77 | 0.24 | 1.14 |
| CD genes | 2.25 | 2.26 | 5.57 | 6.86 | 0.18 | 0.79 | 0.25 | 1.17 |
| HO genes | 1.00 | 1.41 | 4.40 | 5.28 | <0.25 | NA | <1 | NA |
| *ΔHO | -56% |  | -21% |  | NA |  | NA |  |
| #P | 0.27 |  | 0.81 |  | NA |  | NA |  |

\*ΔHO, the % increase of the mean of the HO genes over the mean of the CD genes. #P, the probability that the indicated values for CD and HO oriented genes are equal calculated from the two-tailed Student's t distribution (22) and adjusted for multiple comparisons using the Benjamini-Hochberg procedure (23).
†Only four of the 76 ribosomal genes are in the HO orientation; their CDSs accumulated 4 BPSs and no indels. BPSs, base pair substitutions; indels, insertions and deletions ≤ 4 bps; CDS, coding sequence; SD, standard deviation; CD, codirectional with replication; HO, head-on to replication; NA, not applicable (no mutations).

We note that large insertions, deletions, and rearrangements have been found to be induced at sites of strong replication-transcription conflicts (15, 16) but our sequencing techniques could not detect indels greater than 4 bp.

*E. coli* has 76 genes encoding ribosomal RNAs and proteins; all but 4 of these are in the CD orientation. Of the 86 genes encoding tRNAs, 52 are in the CD orientation (Table S1). Most of these genes are highly expressed; in addition, the tRNA genes are small. Thus, these genes are outliers in expression, size, and orientation and their inclusion in the analysis could bias the results. Therefore, we repeated our analyses with these genes excluded. As shown in Table 3, there was still no significant differences in the frequencies of BPSs or indels between CD and HO oriented CDSs. (See below for analyses of the excluded genes.)

Analyzing the results of mutation accumulation experiments with *B. subtilis*, Schroeder *et al*. (21) found that the slope of the regression line of BPSs/CDS versus CDS length was ∼30% greater for HO-oriented genes than for CD-oriented genes. However, this difference disappeared when the few extreme outliers for length were eliminated. Our results with *E. coli* did not show a difference in the slopes of the regression lines overall (Table S3). However, within the subset of highly expressed genes minus the tRNA genes, to be discussed below, the slope of BPSs/CDS versus CDS length was about 10% greater for HO-oriented genes than for CD-oriented genes (the unadjusted P value for the difference was 0.04 but increased to 0.08 when adjusted for multiple comparisons.)

### Within a subset of highly expressed genes the frequencies of indels, but not of BPSs, are biased toward genes oriented HO to replication

Using our RNA-seq data, we identified a set of highly expressed genes by the criterion that the log_10_(TPM) (transcripts per kilobase per million) for a gene during at least one of the three growth phases (lag, log, and stationary) should be equal to or greater than one standard deviation above the mean log_10_(TPM) for that phase. The result was 770 genes, 483 in the CD orientation and 287 in the HO orientation. As shown in Table 2, among these genes there was no correlation between mutation frequency and expression level. There was also no statistically significant difference in BPSs/CDS between the CD and HO orientations (Table 3). However, among the highly-expressed genes, the frequency of indels/CDS of HO-oriented genes was nearly twice that of CD oriented genes, a difference that was significant at the 5% level when unadjusted but fell to 10% when adjusted for multiple comparisons (Table 3). Dividing the indels/CDS by the length of the CDS increased the bias to nearly three-fold but decreased the significance level to 7% (13% adjusted).

Eliminating the ribosomal and tRNA genes from the subset of highly-expressed genes did not change the BPSs/CDS results but reduced 5 to 10-fold the difference in indels/CDS between CD and HO oriented genes (Table 3). Thus, the bias for indels to be in HO-oriented genes among highly expressed genes is mostly due to the genes that were removed, especially the tRNA genes. We analyze tRNA genes at more length below.

We also identified and analyzed 352 genes that were highly expressed in all three conditions and obtained the same results (Data not shown but available in the IUScholarWorks Repository http://hdl.handle.net/2022/25294.)

### Mutation frequencies in essential genes are not significantly biased by orientation relative to replication

The EcoCyc database (32) lists 358 essential genes for MG1655 *E. coli* strains growing in LB medium. Of these, 252 are oriented CD to replication and 106 are oriented HO to replication. The frequency of BPSs in the CDSs of CD-oriented essential genes was slightly higher than in the CDSs of HO-oriented essential genes, but not significantly so (Table 3). Not surprisingly, since indels tend to be knockout mutations, there were only 31 indels recovered in the CDSs of essential genes and these were distributed 15 in the CD genes and 16 in the HO genes, giving frequencies that appear to be different but that are not statistically significantly different (Table 3).

### Mutations in gene promoters are not biased by orientation relative to replication

The Regulon DB (33) database lists 8568 promoters for MG1655 *E. coli* strains, 4956 of which are oriented CD to replication and 3612 of which are oriented HO to replication. We identified the mutations recovered in our experiments in a region from 60 bp upstream to the transcriptional start sites (TSSs) of these promoters. Neither the frequencies of BPSs nor indels in these promoters differed between the two orientations (Table S4).

To test if a gene’s expression levels affected the frequencies of mutations in its promoter we analyzed the promoters of 3816 genes to which we could assign our RNA-seq values; 2022 of these are oriented CD and 1794 are oriented HO. Among these there was still no difference between the frequencies of BPSs or indels in promoters in the CD versus HO orientation (Table S4). The promoters of the highly expressed genes described above also showed no significant difference between the two orientations either for BPSs or indels (Table S4).

The mutation frequencies in the promoters of essential genes were predictably low, particularly for indels (Table S4). But there was a surprising anomality. No indels were recovered in the promoters of HO-oriented essential genes, but 15 were recovered in the promoters of CD-oriented essential genes. However, 7 of these indels were at a single site, a run of 9 A:T base pairs within the first promoter of the *glnS* gene (positions -35 to -27); an additional 3 indels occurred at a run of 5 G:C base pairs within the second promoter of the *hemA* gene (positions -47 to -43). Thus, this apparent bias in mutation frequency is largely due, instead, to a bias in mutational targets, which will be further discussed below.

As mentioned above, two studies found that a specific BPS, a transition at the 3′T of the −10 element (T_–7_), was induced in promoters in the HO orientation (15, 16). In our collection of 946 A:T transitions in promoters, only 22 were within 20 Nts upstream of a TSS (Fig. S1). Of these only 7 were a transition at T_–7_ of a possible −10 element; two of these were in the CD orientation and 5 were in the HO orientation. While these small numbers would not support the hypothesis that replication-transcription conflict specifically induces a transition at T_–7_, it is possible that our experiments would not detect this mutation. This particular BPS is not elevated in *B. subtilis* MMR^—^ strains, most likely because it is not produced by simple replication errors (15). Thus, it would be swamped out by the uncorrected replication errors in our MMR^—^ strains. In addition, T_–7_ is an important base for promoter recognition and the initiation of transcription (34) and, in *B. subtilis*, this mutation prevents the transcription of many house-keeping genes (J.D. Wang, personal communication). If this is true in *E. coli* as well, transitions of T_–7_ would be selected against in our MA experiments.

### Mutation rates verify the results obtained with mutation frequencies

For the analyses given above we used all the mutations recovered in from 10 mutation accumulation experiments involving 334 independent mutation accumulation lines. In these experiments the lines were propagated for a minimum of 400 generations and a maximum of 3000 generations, giving a total of nearly 250,000 generations (19). These data allow very good estimates of the rate at which mutations accumulated per generation per Nt across the whole genome. Fig. 2 shows that these mutation rates are congruent with the results using mutation frequencies presented above.

**Figure 2.**
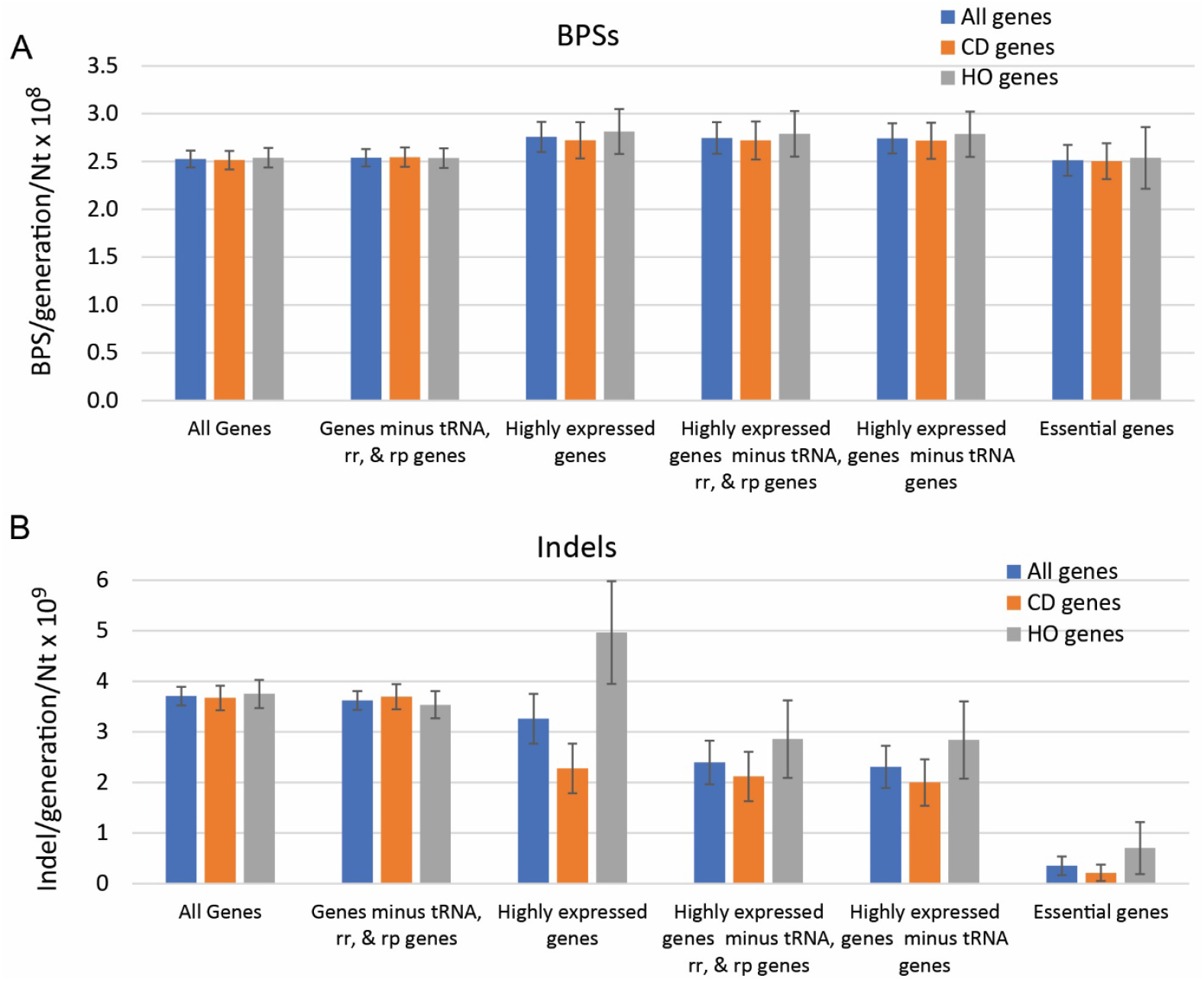
Mutation rates in the coding sequences (CDSs) of all genes and in genes oriented CD and HO to replication. (A) Base pair substitutions (BPSs) per generation per nucleotide x 10^8^ in the CDSs of the indicated genes. (B) Indels (± ≤ 4 bp) per generation per nucleotide x 10^9^ in the CDSs of the indicated genes. Bars represent means and error bars are 95% CLs. Nt, nucleotide; CD, codirectional to replication; HO, head-on to replication.

### Mutation biases in tRNA genes

For their length, the mutation frequencies among tRNA genes were unusually high, particularly for indels; per Nt, tRNAs accumulated twice as many BPSs and nearly 20 times as many indels as did other genes (Table 4). However, examination of the distribution of these mutations reveals that a few of the tRNA genes were heavily mutated while the majority accumulated no or few mutations. Of the 117 indels in tRNA genes, 100 were in just 4 genes, the homologues *leuP, leuQ, leuV*, and *leuT*. Ninety-five of these occurred in the 8 G:C bp run and 3 occurred in the 5 G:C bp run that are in each gene [in the tRNAs, 5 of the 8 Cs encoded by the 8-bp run base pair with the 5 Gs encoded by the other run to form the stem of the TΨC loop (35)].

**Table 4.**
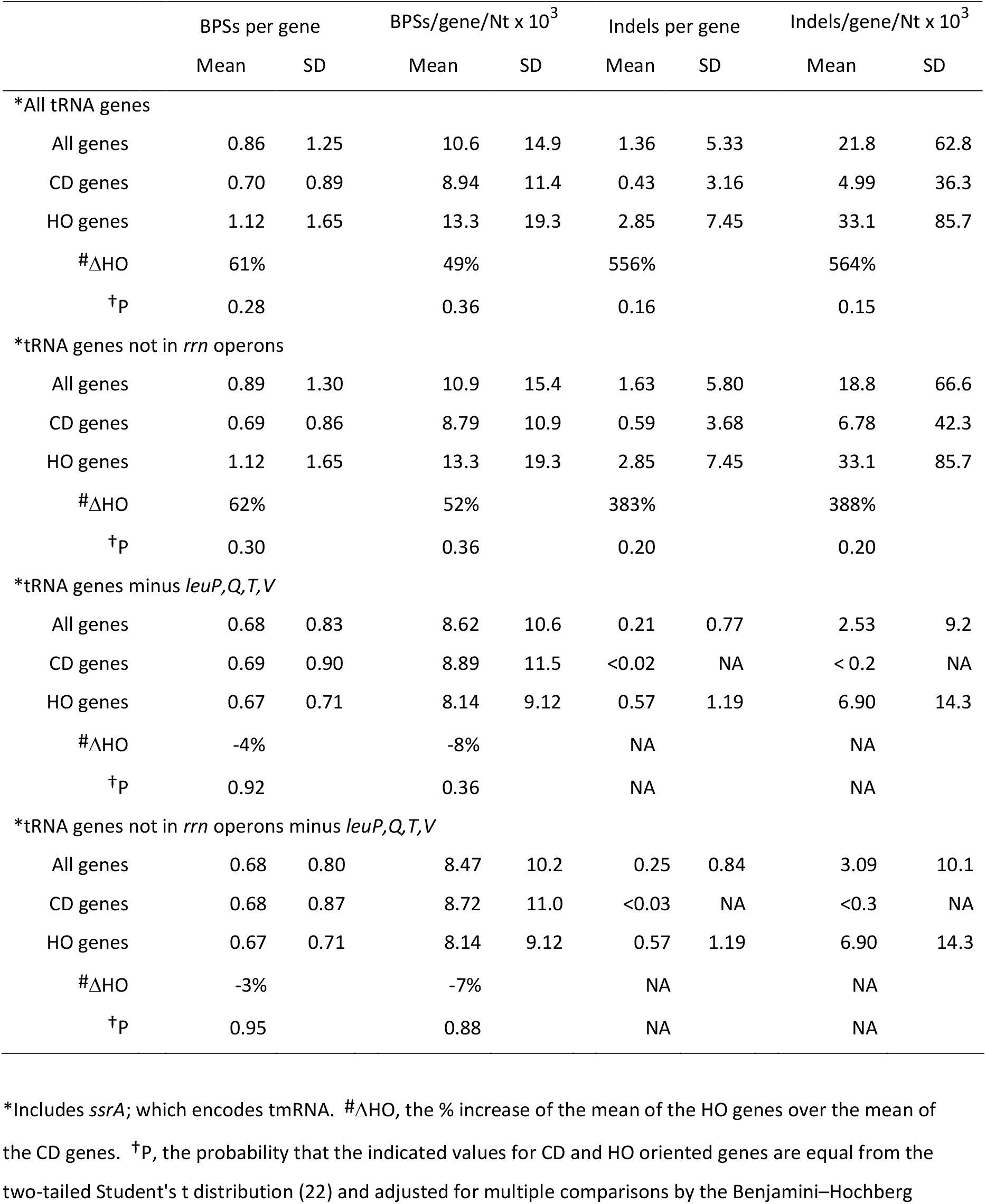
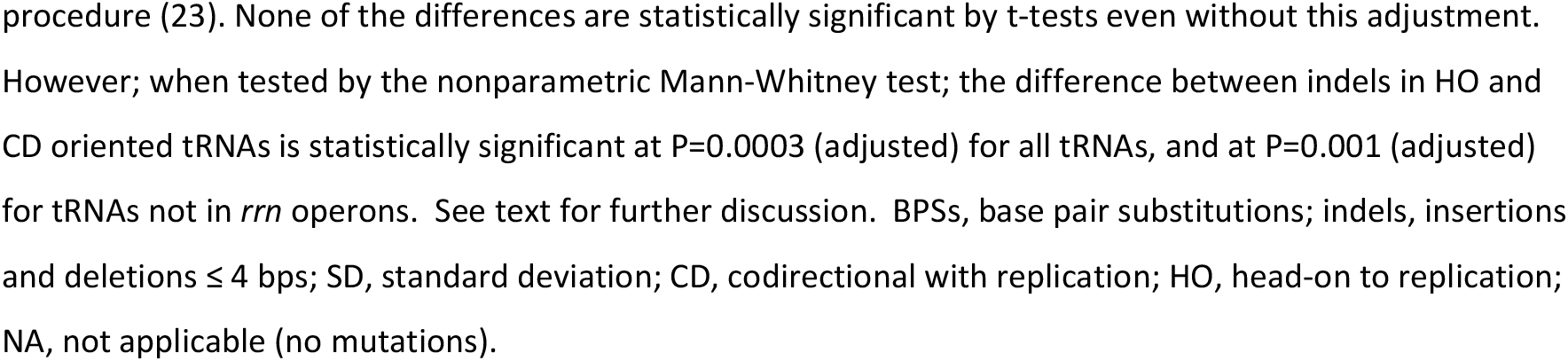
Comparison of the frequencies of mutations in tRNA genes oriented CD versus HO to replication

|  | BPSs per gene |  | BPSs/gene/Nt x 10<sup>3</sup> |  | Indels per gene |  | Indels/gene/Nt x 10<sup>3</sup> |  |
| --- | --- | --- | --- | --- | --- | --- | --- | --- |
|  | Mean | SD | Mean | SD | Mean | SD | Mean | SD |
| *\*All tRNA genes* |  |  |  |  |  |  |  |  |
| All genes | 0.86 | 1.25 | 10.6 | 14.9 | 1.36 | 5.33 | 21.8 | 62.8 |
| CD genes | 0.70 | 0.89 | 8.94 | 11.4 | 0.43 | 3.16 | 4.99 | 36.3 |
| HO genes | 1.12 | 1.65 | 13.3 | 19.3 | 2.85 | 7.45 | 33.1 | 85.7 |
| #ΔHO | 61% |  | 49% |  | 556% |  | 564% |  |
| †p | 0.28 |  | 0.36 |  | 0.16 |  | 0.15 |  |
| *\*tRNA genes not in rrn operons* |  |  |  |  |  |  |  |  |
| All genes | 0.89 | 1.30 | 10.9 | 15.4 | 1.63 | 5.80 | 18.8 | 66.6 |
| CD genes | 0.69 | 0.86 | 8.79 | 10.9 | 0.59 | 3.68 | 6.78 | 42.3 |
| HO genes | 1.12 | 1.65 | 13.3 | 19.3 | 2.85 | 7.45 | 33.1 | 85.7 |
| #ΔHO | 62% |  | 52% |  | 383% |  | 388% |  |
| †p | 0.30 |  | 0.36 |  | 0.20 |  | 0.20 |  |
| *\*tRNA genes minus leuP,Q,T,V* |  |  |  |  |  |  |  |  |
| All genes | 0.68 | 0.83 | 8.62 | 10.6 | 0.21 | 0.77 | 2.53 | 9.2 |
| CD genes | 0.69 | 0.90 | 8.89 | 11.5 | <0.02 | NA | < 0.2 | NA |
| HO genes | 0.67 | 0.71 | 8.14 | 9.12 | 0.57 | 1.19 | 6.90 | 14.3 |
| #ΔHO | -4% |  | -8% |  | NA |  | NA |  |
| †p | 0.92 |  | 0.36 |  | NA |  | NA |  |
| *\*tRNA genes not in rrn operons minus leuP,Q,T,V* |  |  |  |  |  |  |  |  |
| All genes | 0.68 | 0.80 | 8.47 | 10.2 | 0.25 | 0.84 | 3.09 | 10.1 |
| CD genes | 0.68 | 0.87 | 8.72 | 11.0 | <0.03 | NA | <0.3 | NA |
| HO genes | 0.67 | 0.71 | 8.14 | 9.12 | 0.57 | 1.19 | 6.90 | 14.3 |
| #ΔHO | -3% |  | -7% |  | NA |  | NA |  |
| †p | 0.95 |  | 0.88 |  | NA |  | NA |  |

\*Includes *ssrA*; which encodes tmRNA. #ΔHO, the % increase of the mean of the HO genes over the mean of the CD genes. †P, the probability that the indicated values for CD and HO oriented genes are equal from the two-tailed Student's t distribution (22) and adjusted for multiple comparisons by the Benjamini–Hochberg
procedure (23). None of the differences are statistically significant by t-tests even without this adjustment. However; when tested by the nonparametric Mann-Whitney test; the difference between indels in HO and CD oriented tRNAs is statistically significant at $P=0.0003$ (adjusted) for all tRNAs, and at $P=0.001$ (adjusted) for tRNAs not in *rrn* operons. See text for further discussion. BPSs, base pair substitutions; indels, insertions and deletions $\leq 4$ bps; SD, standard deviation; CD, codirectional with replication; HO, head-on to replication; NA, not applicable (no mutations).

Of the 17 indels that occurred in the other tRNA genes, all but 3 occurred in mononucleotide runs ≥ 5 Nts. Because of the strong mutagenic consequence of such runs, we checked to see if tRNA genes have an overabundance of them. As shown in Fig. S2, the number and extent of runs in most tRNA genes do not differ from that for the genome as a whole. However, the 8 G:Cs runs in *leuP, leuQ, leuV*, and *leuT* are outliers. Possibly also outliers are 7 G:Cs runs in *serW* and *serX* (also forming the stem of the TΨC loop) which accounted for 5 and 2 indels, respectively. Of these 6 genes, only *leuT* is oriented CD to replication.

The four homologous tRNA-Leu genes provide an interesting test of whether HO oriented genes have higher mutation rates than CD oriented genes simply because of their orientation relative to replication. *leuP, leuQ*, and *leuV* are in an operon-oriented HO to replication and *leuT* is in a different operon oriented CD to replication. All four genes have similar high expression levels in all three growth stages (Table S2). The numbers of indels recovered in these genes were 23, 29, 25, and 23, respectively, which obviously do not differ statistically. Thus, in this natural experiment in which an identical sequence appears in both CD and HO orientation, the accumulated number of indels/gene was also identical. However, because three of the genes are oriented HO and only one is oriented CD, it mistakenly appears that the HO genes are more highly mutated because of their orientation relative to replication.

The four homologous tRNA-Leu genes also accounted for 18 of the 74 BPSs in tRNA genes. In contrast to the indels, the frequencies of these were biased: *leuP, leuQ*, and *leuV* accumulated 6, 4, and 7 BPSs, but *leuT*, accumulated only 1 BPS. In Supplemental Text S1 and Fig. S3 we explain how this bias may also be due to mutation target placement.

### BPSs are not biased in stress response genes

Merrikh (37) suggested that the placement of stress-response genes in the HO orientation is evolutionarily advantageous. She hypothesized that the accumulation of BPSs in these genes would increase the chance of adaptive mutations, allowing the cell to successfully meet new environmental challenges. Although during our mutation accumulation experiments the cells are not exposed to exogenous stresses, they are exposed to endogenous stresses. For example, in the absence of repair pathways, reactive oxygen species are particularly mutagenic during mutation accumulation experiments (38). To test whether the BPSs that occurred during our mutation accumulation experiments were biased toward HO-oriented stress-response genes, we compared the frequencies of BPSs in the CDSs of CD- and HO-oriented genes that are part of two global stress-response networks, the RpoS-mediated general stress response, and the SOS response to DNA damage.

RpoS is a sigma factor that is induced as cells enter stationary phase and by a variety of other stresses [reviewed in (39)]. Directly or indirectly, it regulates over 600 genes, many of which encode or regulate DNA repair activities. We analyzed the accumulated mutations in the CDSs of 605 genes that are upregulated by RpoS and in a subset of 98 of these that are particularly responsive to RpoS (40). There were no significant differences in BPSs/CDSs in genes oriented CD versus HO to replication (Table 5).

**Table 5.** Comparisons of the frequencies of BPSs in stress-induced CDS oriented CD versus HO to replication

|  |  | No. CDSs | No. BPSs | BPSs per CDS |  | BPSs/CDS/Nt X 10<sup>3</sup> |  |
| --- | --- | --- | --- | --- | --- | --- | --- |
|  | No. CDSs | with BPSs | in CDSs | Mean | SD | Mean | SD |
| All genes upregulated by RpoS |  |  |  |  |  |  |  |
| All genes | 605 | 573 | 4008 | 6.62 | 5.32 | 6.68 | 3.66 |
| CD genes | 349 | 334 | 2231 | 6.39 | 4.85 | 6.93 | 3.63 |
| HO genes | 256 | 239 | 1777 | 6.94 | 5.90 | 6.34 | 3.68 |
| *ΔHO |  |  |  | 9% |  | -9% |  |
| #p |  |  |  | 0.34 |  | 0.10 |  |
| Genes upregulated by low levels of RpoS |  |  |  |  |  |  |  |
| All genes | 98 | 92 | 671 | 6.85 | 5.01 | 7.17 | 3.73 |
| CD genes | 55 | 51 | 382 | 6.95 | 5.30 | 7.46 | 3.67 |
| HO genes | 43 | 41 | 289 | 6.72 | 4.68 | 6.80 | 3.81 |
| *ΔHO |  |  |  | -3% |  | -9% |  |
| #p |  |  |  | 0.88 |  | 0.52 |  |
| †All known or predicted SOS genes |  |  |  |  |  |  |  |
| All genes | 123 | 114 | 780 | 6.34 | 6.14 | 6.51 | 4.09 |
| CD genes | 70 | 65 | 448 | 6.40 | 6.31 | 6.40 | 4.16 |
| HO genes | 53 | 49 | 332 | 6.26 | 5.98 | 6.66 | 4.03 |
| *ΔHO |  |  |  | -2% |  | 4% |  |
| #p |  |  |  | 0.93 |  | 0.83 |  |
| †All genes upregulated by DNA damage |  |  |  |  |  |  |  |
| All genes | 78 | 73 | 472 | 6.05 | 5.92 | 6.48 | 4.29 |
| CD genes | 41 | 39 | 268 | 6.54 | 6.56 | 6.89 | 4.73 |
| HO genes | 37 | 34 | 204 | 5.51 | 5.15 | 6.02 | 3.75 |
| *ΔHO |  |  |  | -16% |  | -13% |  |
| #p |  |  |  | 0.58 |  | 0.51 |  |
| †LexA repressed genes |  |  |  |  |  |  |  |
| All genes | 56 | 52 | 347 | 6.20 | 6.42 | 6.64 | 4.10 |
| CD genes | 28 | 26 | 190 | 6.79 | 7.53 | 6.68 | 4.35 |
| HO genes | 28 | 26 | 157 | 5.61 | 5.15 | 6.60 | 3.91 |
| *ΔHO | | | | -17% | | -1% | |
| #p |  |  |  | 0.63 |  | 0.96 |  |

\* $\Delta$ HO, the % increase of the mean of the HO genes over the mean of the CD genes. #P, the probability that the indicated values for CD and HO oriented genes are equal calculated from the two-tailed Student's t distribution (22) and adjusted for multiple comparisons by the Benjamini–Hochberg procedure (23). <sup>†</sup>The numbers of genes differ from those in Simmons *et al.* (2008) (41) because our strains do not have three damage-induced genes, two of which are on the prophage $\phi$ 14. CDSs, coding sequence; BPSs, base pair substitutions; SD, standard deviation; CD, codirectional with replication; HO, head-on to replication.

DNA damage induces the expression of approximately 80 genes, 57 of which are known to be repressed by the SOS repressor LexA; an additional 44 genes have predicted LexA binding sequences but have not been experimentally shown to be part of the SOS response [reviewed in (41)]. We analyzed the BPSs in the CDSs of all three of these subsets of genes and found no significant differences in mutation frequencies between CDSs oriented CD versus HO to replication (Table 5).

As reported for *B. subtilis* (5), the placement of these stress-response genes in *E. coli* is biased toward the CD orientation (Table 5). Thus, these results provide no support for the hypothesis that stress-response genes are oriented HO to replication so that adaptive mutations will accumulate.

### Transcription-coupled repair does not cause or prevent HO versus CD mutational bias

When RNA polymerase encounters a DNA lesion and stalls, it triggers a rescue and repair pathway called transcription-coupled repair (TRC) (42). In *E. coli* and other bacteria, the Mfd (mutation frequency decline) protein initiates this repair by dislodging the transcription complex and recruiting the nucleotide excision repair pathway to remove the DNA lesion (43). Although the activity of Mfd might be expected to resolve replication-transcription conflicts, several papers have reported that in doing so Mfd creates mutations, at least at certain DNA sites (14, 44, 45).

We previously reported that, as determined by MA-WGS experiments, loss of Mfd had little effect on the mutation rates of MMR-defective strains (19). Deletion of the *mfd* gene in MMR^—^ strains increased the genomic rates of BPSs by about 12% (2.75 vs 2.45 × 10^−8^ BPSs/generation/Nt) and of indels by 28% (5.54 vs 4.33 × 10^−9^ indels/generation/Nt) (Fig. 3). Thus, in contrast to the reports cited above, we found TRC is somewhat antimutagenic. This finding is reminiscent of the original phenotype of Mfd^+^, which is the decline in mutations when *E. coli* strains are briefly incubated without protein synthesis after UV exposure (46).

**Figure 3.**
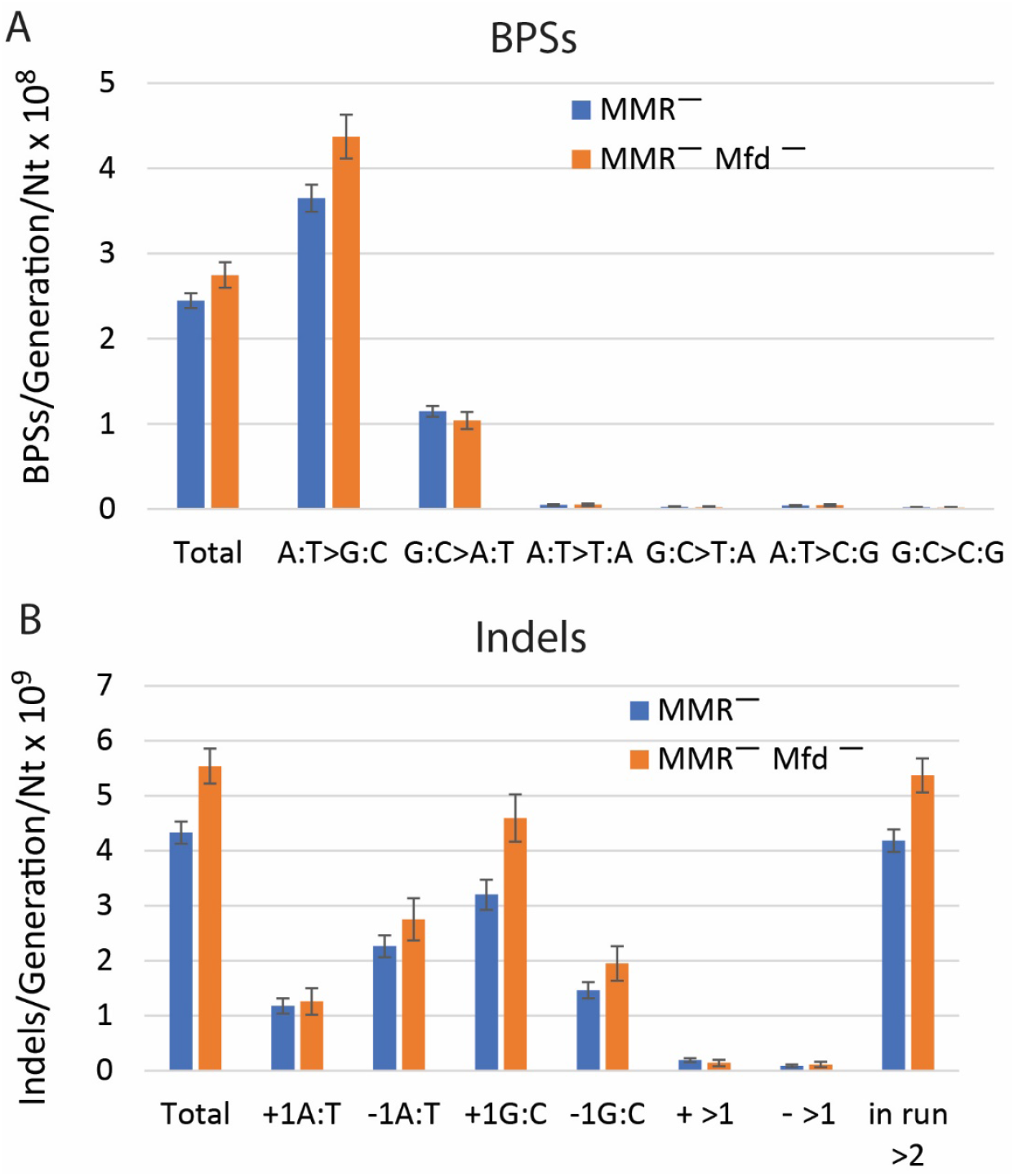
Conditional mutation rates across the genome of *E. coli* strains that are defective for mismatch-repair (MMR^—^) and defective for both mismatch repair and transcription coupled repair (MMR^—^ Mfd^—^). Mutations per generation are normalized by the number of Nts of each type (*e*.*g*., A:T to G:C mutations per generation are divided by the number of A:T bps in the genome). (A) Base pair substitutions (BPSs) per generation per nucleotide x 10^8^. (B) Indels (± ≤ 4 bp) per generation per nucleotide x 10^9^. These results include both CDSs and non-CDSs, so the values for total mutations in the MMR^—^ strains are slightly greater than seen in Figure 2, which include only CDSs. Bars represent means and error bars are 95% CLs. The BPSs data are from Foster *et al*., 2018 (19); the indel data were not previously reported. Nt, nucleotide.

We reanalyzed the data from our MA-WGS experiments to see if loss of Mfd had any impact on the frequency of mutations in CD versus HO oriented genes. Combining our two experiments conducted with Mfd^—^ MMR^—^ strains resulted in 2,582 BPSs and 426 indels in CD oriented genes and 2,265 BPSs and 364 indels in HO oriented genes (Table S1). As shown in Table 6, overall, there were no significant differences in mutation frequencies between genes in the two orientations. There were large apparent increases in the frequencies of both BPSs and indels in HO-oriented versus CD-oriented tRNA genes, but these results are based on only 18 BPSs in 14 genes and 29 indels in 7 genes (see Table S1). And, as with the Mfd^+^ MMR^—^strains, in the Mfd^—^ MMR^—^ strains the HO orientation of *leuP, leuQ, leuV, serW* and *serX* genes accounted for most of the bias: 23 of the 24 indels and 4 of the 11 BPSs in the HO oriented genes were in these genes, and 19 indels and 3 BPSs were in just *leuP, leuQ*, and *leuV*.

**Table 6.** Comparisons of the frequencies of mutations in CDSs oriented CD versus HO to replication in strains defective for both MMR and Mfd

|  | BPSs/CDS |  | BPSs/CDS/Nt X 10<sup>3</sup> |  | Indels/CDS |  | Indels/CDS/Nt X 10<sup>3</sup> |  |
| --- | --- | --- | --- | --- | --- | --- | --- | --- |
|  | Mean | SD | Mean | SD | Mean | SD | Mean | SD |
| All genes |  |  |  |  |  |  |  |  |
| All genes | 1.07 | 1.34 | 1.17 | 1.67 | 0.18 | 0.65 | 0.28 | 2.38 |
| CD genes | 1.05 | 1.34 | 1.15 | 1.58 | 0.17 | 0.56 | 0.24 | 1.55 |
| HO genes | 1.11 | 1.34 | 1.20 | 1.76 | 0.18 | 0.75 | 0.32 | 3.11 |
| *ΔHO | 6% |  | 4% |  | 3% |  | 34% |  |
| #p | 0.21 |  | 0.45 |  | 0.87 |  | 0.40 |  |
| Highly expressed genes |  |  |  |  |  |  |  |  |
| All genes | 0.86 | 1.19 | 1.34 | 2.41 | 0.13 | 0.63 | 0.61 | 5.39 |
| CD genes | 0.84 | 1.17 | 1.31 | 2.20 | 0.09 | 0.48 | 0.31 | 2.87 |
| HO genes | 0.90 | 1.22 | 1.39 | 2.73 | 0.20 | 0.81 | 1.12 | 7.99 |
| *ΔHO | 6% |  | 6% |  | 117% |  | 263% |  |
| #p | 0.70 |  | 0.79 |  | 0.09 |  | 0.18 |  |
| Highly expressed genes minus tRNA genes |  |  |  |  |  |  |  |  |
| All genes | 0.91 | 1.21 | 1.20 | 1.56 | 0.11 | 0.46 | 0.23 | 1.50 |
| CD genes | 0.90 | 1.19 | 1.24 | 1.62 | 0.09 | 0.45 | 0.21 | 1.25 |
| HO genes | 0.94 | 1.25 | 1.13 | 1.46 | 0.14 | 0.47 | 0.26 | 1.85 |
| *ΔHO | 4% |  | -9% |  | 51% |  | 28% |  |
| #p | 0.81 |  | 0.48 |  | 0.33 |  | 0.77 |  |
| †All tRNA genes |  |  |  |  |  |  |  |  |
| All genes | 0.21 | 0.51 | 2.61 | 6.43 | 0.34 | 1.33 | 3.89 | 15.33 |
| CD genes | 0.13 | 0.39 | 1.66 | 5.03 | 0.09 | 0.69 | 1.08 | 7.89 |
| HO genes | 0.33 | 0.65 | 4.12 | 8.05 | 0.73 | 1.92 | 8.39 | 22.12 |
| *ΔHO | 152% |  | 148% |  | 671% |  | 674% |  |
| #p | 0.20 |  | 0.21 |  | 0.15 |  | 0.15 |  |
| †tRNA genes minus *leuP,Q,T,V* |  |  |  |  |  |  |  |  |
| All genes | 0.17 | 0.47 | 2.17 | 6.00 | 0.06 | 0.36 | 0.71 | 4.19 |
| CD genes | 0.12 | 0.38 | 1.48 | 4.89 | <0.02 | NA | <0.25 | NA |
| HO genes | 0.27 | 0.58 | 3.39 | 7.50 | 0.17 | 0.59 | 1.95 | 6.82 |
| *ΔHO | 131% |  | 129% |  | NA |  | NA |  |
| #p | 0.33 |  | 0.34 |  | NA |  | NA |  |

\*ΔHO, the % increase of the mean of the HO genes over the mean of the CD genes. #P, the probability that the indicated values for CD and HO oriented genes are equal calculated from the two tailed Student's t distribution (22) and adjusted for multiple comparisons using the Benjamini-Hochberg procedure (23). By the nonparametric Mann-Whitney test the difference between indels in CDSs oriented HO versus CD is statistically significant at (adjusted) P= 0.04 (Indels/CDS) and P=0.05 (Indels/CDS/Nt) for highly-expressed genes and (adjusted) P= 0.01 for both Indels/tRNA gene and Indels/tRNA gene/Nt. See text for further discussion. <sup>†</sup>Includes *ssrA*; which encodes tmRNA. BPSs, base pair substitutions; indels, insertions and deletions ≤ 4 bps; CDS, coding sequence; SD, standard deviation; CD, codirectional with replication; HO, head-on to replication; NA, not applicable (no mutations).

### Gene orientation does not influence the types of BPSs that occur

Million-Weaver *et al*. (14) reported that induction of a high level of transcription of a *Bacillus subtilis* mutant *hisC* allele increased its reversion rate about 7-fold when the gene was in the HO orientation, but only about 4-fold when the gene was in the CD orientation. About 60% of the induced mutations in the HO-oriented gene were due to DNA PolY1, a member of the Y-family of error-prone polymerases, and about 50% were T to C transitions. T to C mutations are frequent when the fidelities of Y-family polymerase are assayed using single-stranded DNA templates (47, 48). However, the *hisC* target was double-stranded DNA, so the reported T to C transition could have been due to either a G inserted opposite the T or a C inserted opposite the A on the other strand. Since Million-Weaver *et al*. (2015) were assaying on-site reversion of a TAG stop codon, we deduce that the T was on the non-transcribed strand and the A was on the transcribed stand.

To test if a similar mutational signature occurred in our experiments, we compared the frequencies of all six types of BPSs in CDSs in the CD and HO orientations (Table 7). We included all six BPSs because: (i) G:C to A:T transitions might be induced by replication-transcription conflicts but could not have been scored in the assay used by Million-Weaver *et al*. (2015) because G:C to A:T change would create a TAA stop codon; and, (ii) *E. coli*’s Y-family polymerases, Pol IV and Pol V, frequently create transversions (47-52). However, the frequencies of transversions are low, so we grouped them into A:T and G:C transversions for each gene.

**Table 7.**
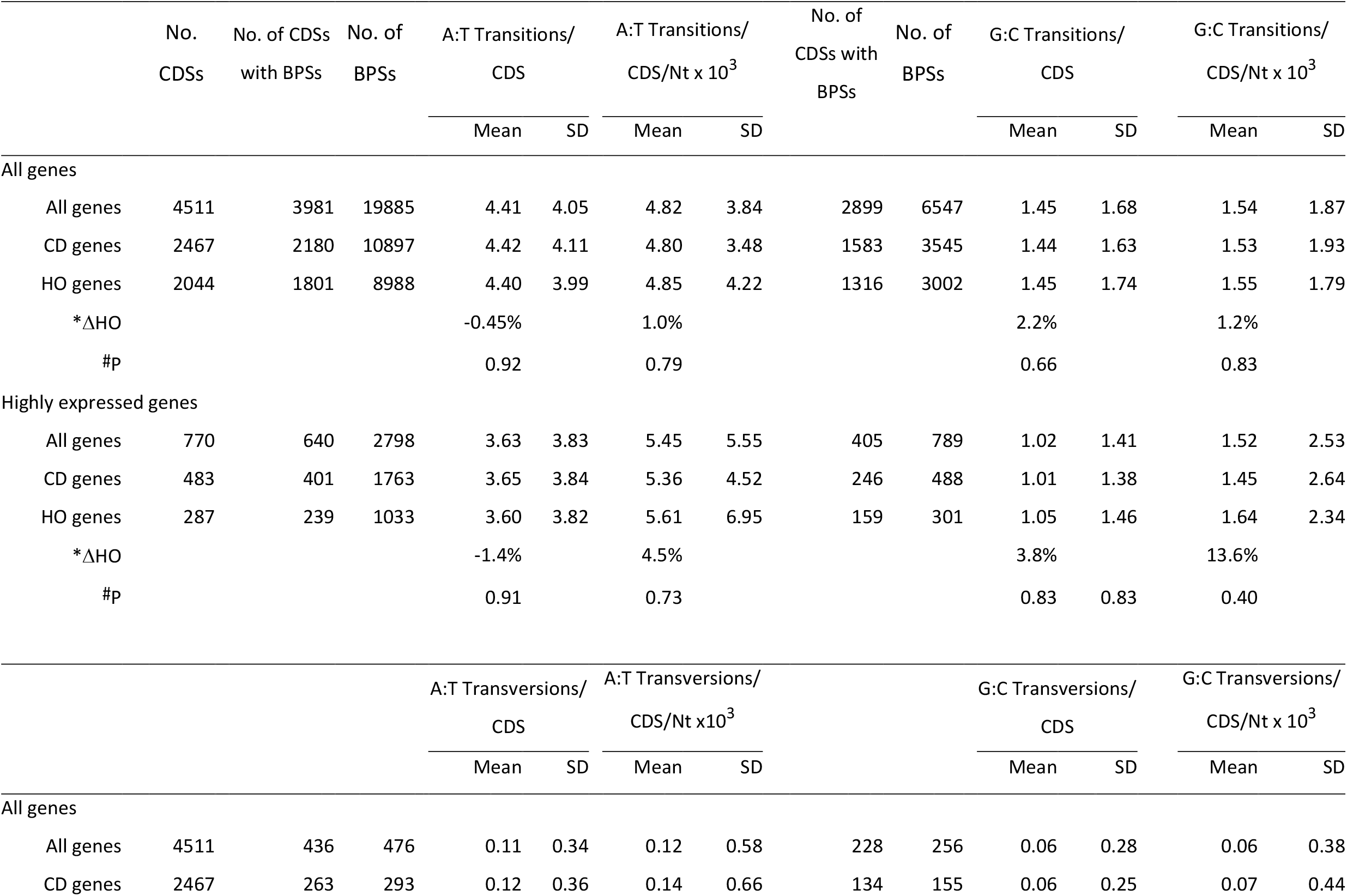
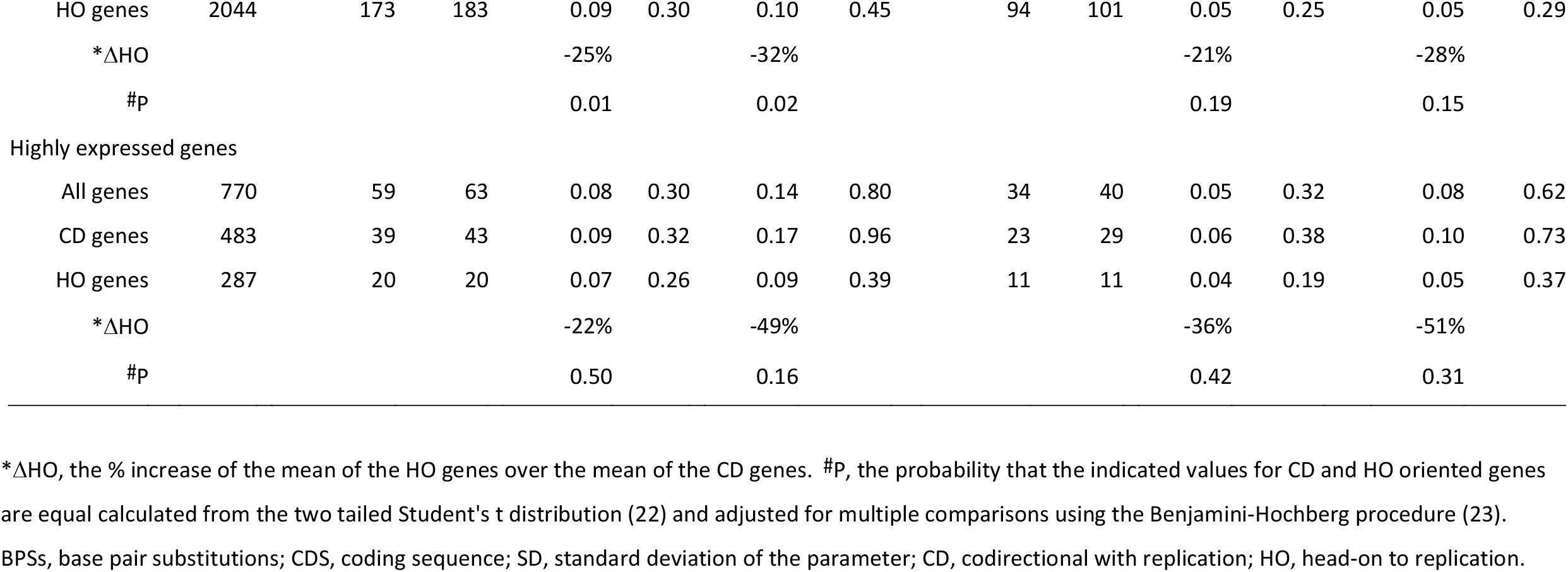
Comparisons of the frequencies of the types of BPSs in CDSs oriented CD versus HO to replication

As shown in Table 7, the frequencies of none of the types of BPSs were enhanced in the HO versus the CD oriented CDSs. Indeed, the only statistically significant difference was a higher mutation frequency among A:T transversions in the CD orientation. There also were no statistically significant orientation biases among the BPSs in the subset of highly expressed genes. Table 8 shows that the frequencies per CDSs of A:T and G:C transitions were not correlated to gene expression levels. (There were too few transversion mutations to make a similar analysis meaningful.)

**Table 8.** Correlations of the frequencies of A:T and G:C transitions per CDS with gene expression levels

|  |  |  |  | log<sub>10</sub>(lag TMP) v. |  | log<sub>10</sub>(log TMP) v. |  | log<sub>10</sub>(stat TMP) v. |  |
| --- | --- | --- | --- | --- | --- | --- | --- | --- | --- |
|  |  |  |  | BPSs/CDS |  | BPSs/CDS |  | BPSs/CDS |  |
|  | *No. of CDSs | No. of CDSs with BPSs | No of BPSs | #ρ | P | #ρ | P | #ρ | P |
| A:T transitions/CDS |  |  |  |  |  |  |  |  |  |
| All genes |  |  |  |  |  |  |  |  |  |
| All genes | 4,495 | 3972 | 19832 | -0.14 | <0.0003 | -0.10 | <0.0003 | -0.15 | <0.0003 |
| CD genes | 2460 | 2176 | 10869 | -0.14 | <0.0003 | -0.11 | <0.0003 | -0.15 | <0.0003 |
| HO genes | 2035 | 1796 | 8963 | -0.14 | <0.0003 | -0.09 | <0.0003 | -0.16 | <0.0003 |
| Highly expressed genes |  |  |  |  |  |  |  |  |  |
| All genes | 770 | 640 | 2,796 | -0.16 | <0.0003 | -0.06 | 0.17 | -0.23 | <0.0003 |
| CD genes | 483 | 401 | 1763 | -0.10 | 0.07 | -0.06 | 0.33 | -0.06 | 0.34 |
| HO genes | 287 | 239 | 1033 | -0.10 | 0.17 | -0.08 | 0.29 | -0.16 | 0.01 |
| G:C transitions/CDS |  |  |  |  |  |  |  |  |  |
| All genes |  |  |  |  |  |  |  |  |  |
| All genes | 4,495 | 2890 | 6532 | -0.16 | <0.0003 | -0.12 | <0.0003 | -0.16 | <0.0003 |
| CD genes | 2460 | 1579 | 3539 | -0.16 | <0.0003 | -0.14 | <0.0003 | -0.17 | <0.0003 |
| HO genes | 2035 | 1311 | 2993 | -0.16 | <0.0003 | -0.11 | <0.0003 | -0.14 | <0.0003 |
| Highly expressed genes |  |  |  |  |  |  |  |  |  |
| All genes | 770 | 405 | 789 | -0.18 | <0.0003 | -0.08 | 0.05 | -0.14 | <0.0003 |
| CD genes | 483 | 246 | 488 | -0.16 | 0.001 | -0.10 | 0.05 | -0.16 | 0.003 |
| HO genes | 287 | 159 | 301 | -0.21 | 0.001 | -0.05 | 0.54 | -0.11 | 0.12 |

\*The numbers of CDSs are smaller than for previous analyses because we did not have RNA-seq data for 16 genes. (See Material and Methods.) # $\rho$ , Pearson's correlation coefficient. Calculating Spearman's rank correlations did not improve the correlations. P, the probability that the correlation occurred by chance calculated from the two-tailed Student's t-distribution (22) and adjusted for multiple comparisons by the Benjamini–Hochberg procedure (23). Note: weak correlations can be statistically significant. Lag, log, and stat, the lag, log (exponential), and stationary phases of growth; TMP, transcripts per kilobase million; CDSs, coding sequences; BPSs, base pair substitutions; CD, codirectional with replication; HO, head-on to replication.

We also checked to see if it mattered which base was on the transcribed strand. As shown in Fig. 4, A:T transitions were almost twice as frequent in HO versus CD oriented CDSs when A was on the transcribed strand, and G:C transitions were about 60% more frequent in HO versus CD oriented CDSs when C was on the transcribed strand. However, this orientation bias may have nothing to do with transcription but, instead, may reflect on which DNA strand the base is located. The transcribed strand in HO oriented CDSs is the lagging-strand template (LGST) and in CD oriented CDSs is the leading-strand template (LDST). As we previously reported (18, 54) across the chromosome A:T and G:C transitions occur about twice as frequently when the A and the C are on the LGST than when these bases are on the LDST. As shown in Fig. 4, this strand bias is sufficient to explain the apparent differences between CD and HO oriented CDSs.

**Figure 4.**
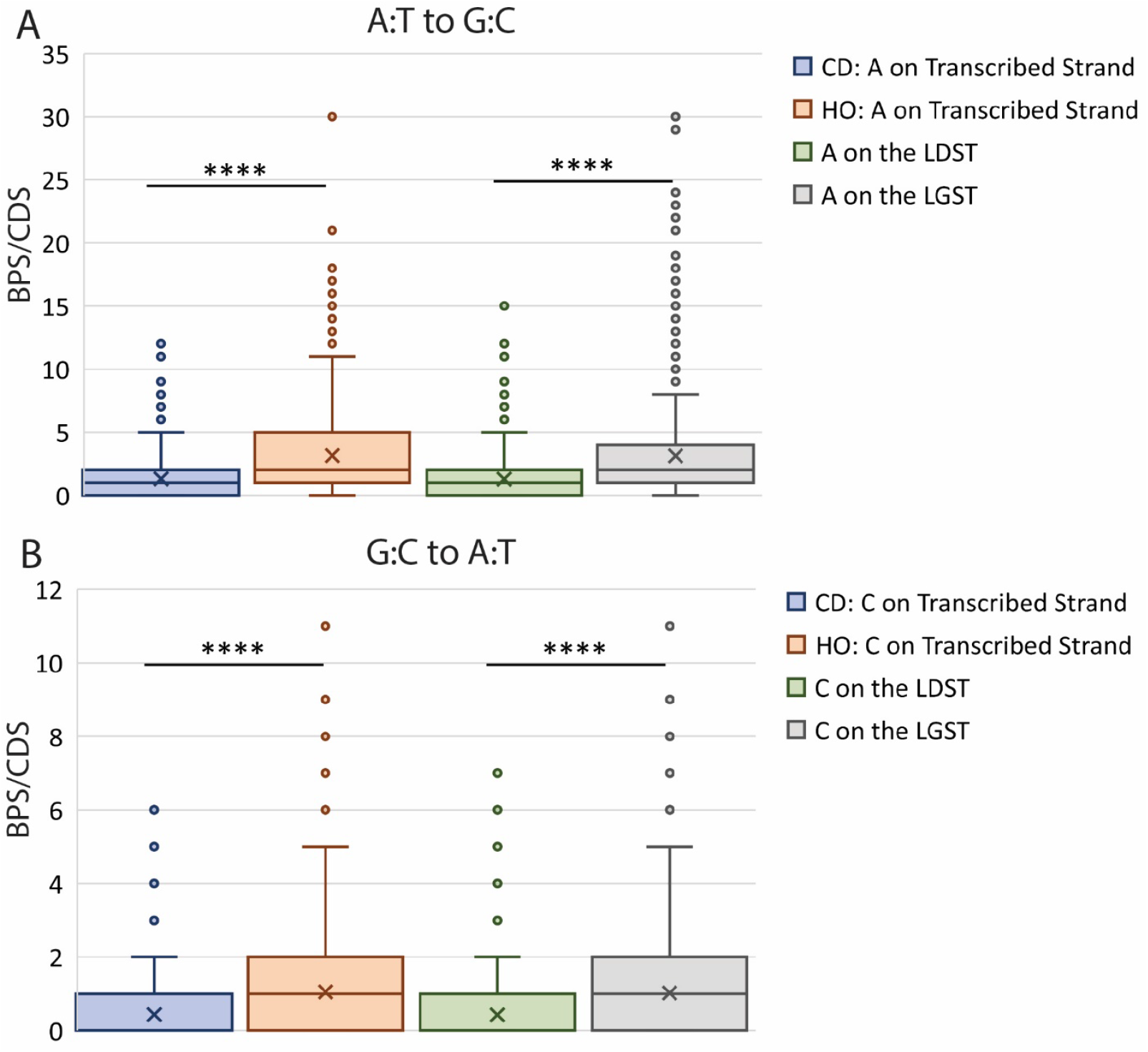
The frequencies of transitions relative to gene orientation and to DNA strand location. (A) The frequencies of A:T to G:C transitions per coding sequence (CDS). (B) The frequencies of G:C to A:T transitions per CDS. Because of the large data variances, the results are presented in both parametric (means) and nonparametric (medians and distributions) form. The X’s in the boxes indicate the means, the horizontal lines in the boxes indicate the medians, the bottoms and tops of the boxes indicate the first (Q1) and third (Q3) quartiles, the “whiskers” indicate the minimum and maximum values that are not outliers, with an outlier defined as equal to or greater than 1.5 times the interquartile range (Q3 minus Q1) below or above Q1 and Q3 (53). The points above the whiskers are the outliers; there are no outliers below the whiskers. For the G:C to A:T results, the medians of the blue and green boxes are 0. The horizontal lines with asterisks indicate that the differences between the indicated means (by Student t tests) or the distributions about the indicated medians (by Mann-Whitney tests) are highly significant (<<0.0003). No other differences are significant. BPSs, base-pair substitution; Nt, nucleotide; CD, codirectional to replication; HO, head-on to replication; LDST, leading strand template; LGST, lagging strand template.

### Comparison of results with *Bacillus subtilis* to those with *E. coli*

While only about half of the genes in *E. coli* are oriented CD to replication, three quarters of the *Bacillus subtilis* genes are so oriented (12). In MS-WGS experiments with MMR-defective strains of *B. subtilis*, Schroeder *et al*.(21) found that HO-oriented genes did have higher rates of BPSs than CD-oriented genes, but this bias was entirely due to bias in the number of the triplet nucleotide sequences that are transition hotspots. Schroeder *et al* (2016) used MMR-defective derivatives of the domesticated *B. subtilis* strain, PY79 (21), whereas we have data from four MA-WGS experiments with MMR-defective derivatives of the non-domesticated *B. subtilis* strain, NCIB 3610 (26). These four strains differ in their competence phenotypes but not in their mutagenic phenotypes (26). Combining the results yielded 9,587 BPSs and 2,100 indels which we reanalyzed and compared to the results reported in Schroeder *et al*. (2016) (see Table S5 for the data discussed below).

As with *E. coli*, the numbers of BPSs/CDS in our *B. subtilis* strains were significantly correlated with CDS length. While BPSs/CDS did not differ significantly between genes in the two orientations, the frequency of BPSs/CDS/Nt was 6% greater in genes oriented HO versus CD to replication. We confirmed the finding of Schroeder *et al*. (2016) that the slope of the regression line of BPSs/CDS versus CDS length of HO oriented genes was greater than that of CD oriented genes, but in our case the slopes differed by only 10% (Table S5B), whereas the difference found by Schroeder *et al*. (2016) appears to be about 30% (but was zero when outliers for length were eliminated). Comparing mutation rates, *i*.*e*., BPSs/generation/Nt across the whole genome (Fig. 5A), the rate in HO oriented genes was 9% greater than in CD oriented genes (P=0.04), a difference that was due to higher rates of G:C to A:T transitions in HO oriented genes, as previously reported (21).

**Figure 5.**
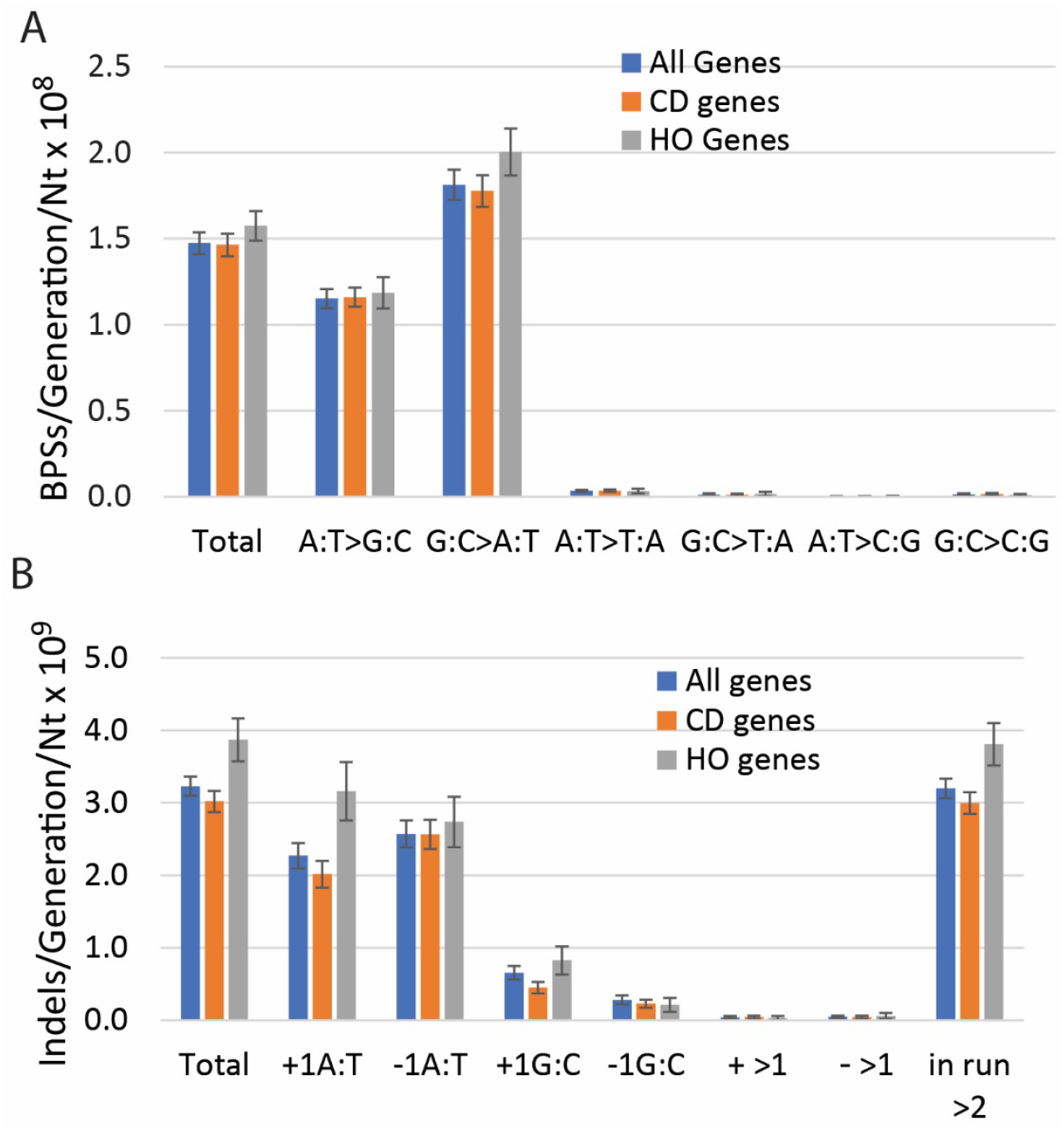
Conditional mutation rates in all the CDSs of genes and in genes oriented CD and HO to replication in *B. subtilis*. Combined results of four MA-WGS experiments with *B subtilis* MMR-defective strains. Mutations per generation are normalized by the number of Nts of each type (*e*.*g*., A:T to G:C mutations per generation are divided by the number of A:T bps in the CDSs). (A) Base pair substitutions (BPSs) per generation per nucleotide x 10^8^. (B) Indels (± ≤ 4 bp) per generation per nucleotide x 10^9^. Bars represent means and error bars are 95% CLs. BPS, base pair substitution, CD, codirection to replication; HO, head-on to replication.

In our strains the frequency of indels/CDS was 18% higher in HO oriented genes than in CD oriented genes; this difference increased to 61% when normalized for CDS length. Comparing indels/generation/Nt across the whole genome (Fig. 5B), genes-oriented HO to replication had a 28% higher rate than did genes oriented CD to replication (P <0.0003). This difference was mostly due to an increase in insertions of A:T bps in mononucleotide runs of A:T bps. In contrast to the results for *E. coli* described above, all but two tRNA genes in *B. subtilis* are oriented CD and so yield little information about mutational bias.

## DISCUSSION

The results presented here provide little evidence that conflicts between DNA replication and transcription are mutagenic in *E. coli* growing under stress-free conditions. Whether considering all genes or various subsets of them, the frequencies of BPSs and indels were not significantly greater in the CDSs or promoters of genes transcribed HO to replication than in genes transcribed CD to replication. The single exception was a greater frequency of indels in HO oriented genes in a subset of highly expressed genes dominated by the tRNA genes. However, the apparent greater bias in tRNA genes was almost entirely due to four homologous genes carrying strong hotspots for indels, three of which are oriented HO and one of which is oriented CD. Overall, this and other studies with a variety of bacterial species support the hypothesis that, in the absence of exogenous stress, the major determinant of spontaneous mutation at a given site is not a conflict between replication and transcription, but, instead, the local sequence context (18, 19, 21, 38, 55, 56).

The hypothesis that conflicts between replication and transcription are mutagenic derives largely from comparisons of mutation rates in reporter genes that were inserted into the *B. subtilis* chromosome in forward and reverse orientations. The mutations scored were reversions of point mutations in genes for amino acid synthesis (13, 14) and inactivation of a gene that makes the bacteria sensitive to trimethoprim (15). These experiments showed that when the gene was oriented HO to replication it had about twice the mutation rate as when it was oriented CD to replication, but only when transcription was artificially induced. A similar result was seen when one fourth of the *B. subtilis* chromosome was inverted, which flipped the normally highly expressed *rpoB* gene to the HO orientation: as a result, the rate of BPSs in *rpoB* conferring resistance to rifampicin increased about 3-fold relative to the normal strain (8).

These artificial constructs maximize replication-transcription conflicts by positioning highly transcribed genes HO to replication. The conflicts then lead to mutations by one mechanism or another. But the mutations that result, such as BPSs and indels that inactivated genes (15), are detrimental and would not be adaptive when not under selection. Unperturbed genomes have evolved to minimize conflict that would be deleterious (because most mutations are deleterious) and so replication-transcription conflicts normally do not contribute to spontaneous mutation rates (J.D. Wang, personal communication).

In addition, there are other mechanisms that can account for an apparent orientation bias in mutation rates. Flipping the orientation of a gene also changes whether its CDS is replicated as the leading or the lagging DNA strand, and the two processes appear to have different error rates. Using reversion of four alleles carrying point mutations in the *lacZ* gene in *E. coli*, Fijalkowska *et al*. (57) found 2 to 4-fold differences in mutation frequencies in one orientation versus the other. Importantly, the higher mutation frequencies were not coordinated with the direction of transcription. When mutations across the entire *lacI* gene were scored in the two orientations, the mutation frequencies were the same, thus not influenced by transcription, but the spectra of mutations were different (58). As we previously reported with a smaller data set (18) and shown here in Fig. 4, in the spectrum of BPSs across the *E. coli* chromosome is strand-specific: A:T to G:C transitions are twice as likely to occur with A templating the lagging strand and T templating the leading strand than in the opposite orientation, whereas G:C to A:T transitions are twice as likely to occur with C templating the lagging strand and G templating the leading strand than in the opposite orientation. This is the same bias found in the experiments with *lacI* (58). Thus, as previously noted (5), when a gene is inverted the DNA context met by the replication machinery for each strand is different, and this fact could result in the mutational biases reported. These biases would be particularly strong when only one or a few specific mutations are scored, as in the reversion assays referenced above.

Given that replication-transcription conflicts are detrimental to genomic stability (6-8, 10), it has been argued that the persistence of HO-oriented genes in genomes means that mutation of these genes must convey an evolutionary advantage (37). Merrikh and colleagues (13, 37, 59) have argued that stress-response genes are oriented HO to replication to increase the probability that these genes will acquire adaptive mutations. But the data presented here for *E. coli* (Table 5) and previously for *B. subtilis* (5) show no evidence that stress-induced genes are preferentially in the HO-orientation. In addition, we found that HO-oriented stress-response genes also are not preferentially mutated (Table 5). Further, Schroeder *et al*. (2020) have convincingly argued that the occurrence of HO-oriented genes in genomes is the result of a balance between inversions that create them and purifying selection that removes them. They further note that, at least in *B. subtilis*, genes oriented HO tend to be non-essential and thus under relaxed selection, which would further account for their persistence as well as for their greater rate of non-synonymous BPSs. Supporting this hypothesis, more than twice as many essential genes in *E. coli* are oriented CD rather than HO to replication (Table S1), and there is no difference in the mutation frequencies between the genes in the two orientations (Table 3).

As mentioned above, the most important determinant of mutation rate at a given site is the local sequence context. For BPSs, the identities of the adjacent bases can influence the mutation rate of a base as much as 400-fold (21). The mutation rates of indels are an exponential function of the length of the homopolymeric run in which they appear (18). Thus, as previously noted (21), apparent differences in mutation rates of genes oriented CD versus HO to replication can be explained to a great extent by differences in the sequence contexts in the two orientations. In addition, our finding that the orientation bias of mutation frequencies in tRNA genes was simply due to gene placement demonstrates that the distribution of specific mutational targets across the genome strongly influences mutation rates. These factors should be evaluated before concluding that a given bias in mutation rates is due to replication-transcription conflicts.

## MATERIALS AND METHODS

### Strains and procedures

All the *E. coli* strains reported in this paper are derivatives of PFM2, a prototrophic *rpoS*^+^ derivative of MG1655 (38). The *B. subtilis* strains are derivatives of the non-domesticated strain NCIB 3610 (60) and were kindly supplied by D. Kearns and M. Konkol. The methods of strain construction, the conditions of growth, and the MA procedures have been published (19, 26).

### RNA sequencing

The RNA-seq procedures are published (26). However, there was an inadvertent error in the description of the samples. Due to a problem in sequencing, one replicate of the samples taken during the lag and stationary phases of growth was lost, so the means of only two replicates for those samples were used. These values are given in Table S2

### Mutation Annotation

For our original mutation calls we used the *E. coli* reference sequence, NC_000913.2 (Version 2) (18). The current reference sequence, NC_000913.3 (Version 3), differs in both the length of the genome (due to insertion sequences) and gene annotation (32). Twenty places where we called mutations in non-coding regions have been updated to genes in Table S2. Our original mutation calls were in protein-coding genes only, but mutations in known RNA genes have also been updated in Table S2. Version 3 was used for our RNA-seq analysis, but we obtained no expression data for 16 genes. Also 146 pseudogenes have been identified in Version 3 for which we do not have mutational data. These minor discrepancies do not change any of the conclusions of this paper.

### Statistical Analyses

Data were fit to Guassian curves using the Mathworks MatLab Curve Fitting app. Statistical tests were computed using either Microsoft Excel or MatLab; the underlying methods are found in reference (22). While in almost all cases the probabilities from parametric tests (Pearson’s linear correlation, Student’s t-tests, F tests) are presented, nonparametric tests were always also used (Spearman’s rank correlation, Mann-Whitney tests). If these probabilities differed, the difference is noted in the text or the Table and Figure legends.

There were over 400 comparisons statistically evaluated for these studies. To compensate for multiple comparisons, we applied the Benjamini-Hochberg correction (23) to the original calculated values. To allow the uncorrected probabilities to be extracted, the relationship between the two probability values is shown in Fig. S4.

For the mutational analysis presented in this paper, the results of 10 MA experiments with MMR^—^ strains, comprising 334 independent MA lines, were treated as one data set. The reasoning behind this treatment was given in reference (19). Briefly, there were no significant differences in the mutation rates or spectra among the experiments, as is clear from the small error bars in Figs. 2A and 2B. The data are dominated by one experiment with a Δ*mutH* strain that ran for nearly 100,000 generations (due to a miscommunication), whereas the rest of the experiments averaged about 20,000 generations. But the results from this experiment are the same as those from the other experiments. Indeed, when the numbers of mutations from each experiment are simply normalized by the generations for that experiment, the means ± SDs are 0.115 ± 0.017 for BPSs and 0.022 ± 0.0014 for indels.

### Data Availability

The sequences and SNPs reported in this paper have been deposited with the National Center for Biotechnology Information Sequence Read Archive https://trace.ncbi.nlm.nih.gov/Traces/sra/ (accession no. SRP013707) and in the IUScholarWorks Repository (http://hdl.handle.net/2022/25294).

## ACKNOWLEDGEMENTS

This research was supported by US Army Research Office Multidisciplinary University Research Initiative (MURI) Award W911NF-09-1-0444 to P.L.F. and H.T., and the National Institutes of Health T32 GM007757 to B.A.N. The funders had no role in study design, data collection and interpretation, or the decision to submit the work for publication. We particularly appreciate E. Popodi, J. Townes, W. Mohammed Ismail, and H. Tang for their past intellectual and technical contributions to this research. We thank E. A Housworth for statistical advice. We thank the following former members of the P.L.F laboratory for technical assistance: H. Bedwell, C. Coplen, M. Durham, J. Eagan, J. Ferlmann, N. Gruenhagen, T. Gruenhagen, J.A. Healy, N. Ivers, C. Klineman, I. Rameses, S. Riffert, H. Rivera, D. Simon, K. Smith, L. Tran, and L. Whitson. Bacterial strains were kindly provided by D. Kearns, M. Konkol, and The National BioResource Project at the (Japanese) National Institute of Genetics. Finally, we thank the reviewers of this paper for useful and insightful comments.

## SUPPLEMENTAL FILES

### SUPPLEMENTAL TABLES

**Table S1. Aggregate gene and mutation data**.

**Table S2. *E. coli* data by gene**

**Table S3. Comparisons of the slopes of BPSs per CDS vs CDS length in Nts between genes oriented CD versus HO to replication**

**Table S4. Comparisons of the frequencies of mutations in the promoters of genes oriented CD versus HO to replication**

**Table S5. *Bacillus subtilis* mutational data and results**

### SUPPLEMENTAL FIGURES

**Figure S1. Search for A:T transitions at the 3′ base in -10 elements**

**Figure S2. The distribution of mononucleotide runs in tRNA genes and across the genome**. long, but

**Figure S3. The effects of differences in orientation on possible BPSs due to slipped-mispairing at runs**.

**Figure S4. The relationship between the calculated probability values for the statistical tests performed for this paper and the Bonferroni-Holm adjusted probabilities values**

### SUPPLEMENTAL TEXT

**Text S1: Base-pair substitution biases in tRNA genes could be due to biased target positions**.

