## Supplemental Files for "DNA Replication-transcription conflicts do not significantly contribute to spontaneous mutations due to replication errors in *Escherichia coli*."

**Text S1**: Base-pair substitution biases in tRNA genes could be due to biased target positions

The four tRNA *leu* homologues accounted for 18 of the 74 BPSs that accumulated in tRNA genes, which was 6-fold greater than the average frequency in the other 82 tRNA genes (Table S1). In contrast to the indels, the frequencies of these 18 BPSs were biased: *leuP*, *leuQ*, and *leuV*, the three genes-oriented HO to replication, accumulated 6, 4, and 7 BPSs, but *leuT*, the gene oriented CD to replication, accumulated only 1 BPS. This result suggests that there is an influence of transcription direction on the production of BPSs. However, consideration of the types of BPSs suggests an alternative explanation. Twelve of the BPSs, 4 in *leuP*, 3 in *leuQ,* and 5 in *leuV*, were transitions at the A:T bp adjacent to the run of 8 G:C bps (Figure S3A ). BPSs adjacent to runs can be ascribed to loop-out of the primer during DNA synthesis, synthesis of an extra base, then realignment to the template, creating a mispair with the base 3′ to the run (1, 2). This could occur during leading strand replication in *leuP*, *Q*, and *V*, but only during lagging strand replication in *leuT*. If we assume that primer loop-out is much more likely during leading strand replication than during lagging strand replication, the mutational bias is explained. Supporting this hypothesis, the one BPS in *leuT*, an A:T to C:G transversion that occurred at the A:T bp adjacent to the 5 G:C bp run could be made during leading strand replication in *leuT*, but did not occur in *leuP*, *Q*, or *V*, where it could only be made during lagging strand replication. (Figure S3B)

One addition note about these special mutations. One transition of the terminal G:C of the 8 G:C bp run occurred in *leuP*. This mutation could be made by loop-out of the template during lagging strand replication in *leuP*, *Q*, and *V*, but only during leading strand replication in *leuT* (Figure S3C). So, it is possible that primer loop-out is more frequent during leading strand replication and template loop-out is more frequent during lagging strand replication.

**Table S1**. Aggregate gene and mutation data

| | No. of CDSs | *E. coli* MMR⁻strains | | | | *E. coli* MMR⁻ Mfd⁻ strains | | | |
|---|---|---|---|---|---|---|---|---|---|
| | | No. of CDS with BPSs | No. of BPSs in CDSs | No. of CDSs with indels | No. of Indels in CDSs | No. of CDS with BPSs | No. of BPSs in CDSs | No. of CDSs with indels | No. of Indels in CDSs |
| All Genes | | | | | | | | | |
| All CDSs | 4,511 | 4,133 | 27,164 | 1,497 | 3,841 | 2,576 | 4,846 | 530 | 790 |
| CD CDSs | 2,467 | 2,261 | 14890 | 810 | 2119 | 1409 | 2,581 | 302 | 426 |
| HO CDSs | 2,044 | 1,872 | 12274 | 687 | 1722 | 1,167 | 2265 | 228 | 364 |
| Gene minus tRNA and ribosomal genes | | | | | | | | | |
| All CDSs | 4,348 | 4035 | 26923 | 1478 | 3711 | | | | |
| CD CDSs | 2,342 | 2185 | 14691 | 803 | 2083 | | | | |
| HO CDSs | 2006 | 1850 | 12232 | 675 | 1628 | | | | |
| Highly expressed genes | | | | | | | | | |
| All CDSs | 770 | 674 | 3688 | 160 | 418 | 381 | 665 | 58 | 103 |
| CD CDSs | 483 | 420 | 2323 | 88 | 195 | 241 | 408 | 28 | 45 |
| HO CDSs | 287 | 254 | 1365 | 72 | 223 | 140 | 257 | 30 | 58 |
| Highly expressed genes minus tRNA and ribosomal genes | | | | | | | | | |
| All CDSs | 657 | 594 | 3486 | 148 | 303 | | | | |
| CD CDSs | 396 | 357 | 2156 | 86 | 171 | | | | |
| HO CDSs | 261 | 237 | 1330 | 62 | 132 | | | | |
| Highly expressed genes minus tRNA genes | | | | | | | | | |
| All CDSs | 713 | 643 | 3633 | 149 | 304 | 370 | 651 | 53 | 76 |
| CD CDSs | 447 | 403 | 2298 | 87 | 172 | 236 | 402 | 27 | 40 |
| HO CDSs | 266 | 240 | 1335 | 62 | 132 | 134 | 249 | 26 | 36 |
| Essential genes | | | | | | | | | |
| All CDSs | 358 | 338 | 2232 | 17 | 31 | | | | |
| CD CDSs | 252 | 242 | 1605 | 11 | 15 | | | | |
| HO CDSs | 106 | 96 | 627 | 6 | 16 | | | | |
| Ribosomal genes | | | | | | | | | |
| All CDSs | 76 | 53 | 166 | 6 | 13 | | | | |
| CD CDSs | 72 | 51 | 162 | 6 | 13 | | | | |
| HO CDSs | 4 | 2 | 4 | 0 | 0 | | | | |

| | | | | | | | | | |
|---|---|---|---|---|---|---|---|---|---|
| *tRNA genes | | | | | | | | | |
| All genes | 86 | 44 | 74 | 13 | 117 | 14 | 18 | 7 | 29 |
| CD genes | 53 | 25 | 37 | 1 | 23 | 6 | 7 | 1 | 5 |
| HO genes | 33 | 19 | 37 | 12 | 94 | 8 | 11 | 6 | 24 |
| *tRNA genes not in *rrn* operons | | | | | | | | | |
| All genes | 72 | 38 | 64 | 13 | 117 | | | | |
| CD genes | 39 | 19 | 27 | 1 | 23 | | | | |
| HO genes | 33 | 19 | 37 | 12 | 94 | | | | |
| *tRNA genes minus *leuP,Q,T,V* | | | | | | | | | |
| All genes | 82 | 40 | 56 | 9 | 17 | 11 | 14 | 3 | 5 |
| CD genes | 52 | 24 | 36 | 0 | 0 | 5 | 6 | 1 | 1 |
| HO genes | 30 | 16 | 20 | 9 | 17 | 6 | 8 | 3 | 5 |
| *tRNA genes not in *rrn* operons minus *leuP,Q,T,V* | | | | | | | | | |
| All genes | 68 | 34 | 46 | 9 | 17 | | | | |
| CD genes | 38 | 18 | 26 | 0 | 0 | | | | |
| HO genes | 30 | 16 | 20 | 9 | 17 | | | | |
| All promoters | | | | | | | | | |
| All genes | 8568 | 1050 | 1235 | 150 | 302 | | | | |
| CD genes | 4956 | 605 | 709 | 84 | 164 | | | | |
| HO genes | 3612 | 445 | 526 | 66 | 138 | | | | |
| Promoters of known genes | | | | | | | | | |
| All genes | 3816 | 477 | 545 | 90 | 167 | | | | |
| CD genes | 2022 | 251 | 288 | 46 | 80 | | | | |
| HO genes | 1794 | 226 | 257 | 44 | 87 | | | | |
| Promoters of highly Expressed Genes | | | | | | | | | |
| All genes | 832 | 101 | 115 | 17 | 38 | | | | |
| CD genes | 469 | 55 | 64 | 8 | 16 | | | | |
| HO genes | 363 | 46 | 51 | 9 | 22 | | | | |
| Promoters of essential genes | | | | | | | | | |
| All genes | 289 | 33 | 39 | 6 | 15 | | | | |
| CD genes | 190 | 22 | 24 | 6 | 15 | | | | |
| HO genes | 99 | 11 | 15 | 0 | 0 | | | | |

*Includes *ssrA*, which encodes tmRNA

MMR = mismatch repair, Mfd = mutation frequency decline, CDSs = coding sequences, BPSs = base pair substitutions, Indels = insertions and deletions ≤ 4 bp, CD = codirectional with replication, HO = head-on to replication.

**Table S3**. Comparisons of the slopes of BPSs per CDS vs CDS length in Nts between genes oriented CD versus HO to replication

| | Slope | SE Slope | intercept | SE intercept | $R^2$ | $*P_F$ |
|---|---|---|---|---|---|---|
| All genes | | | | | | |
| All genes | 0.0066 | 0.0001 | -0.06 | 0.08 | 0.65 | <0.0003 |
| CD genes | 0.0066 | 0.0001 | -0.07 | 0.11 | 0.65 | <0.0003 |
| HO genes | 0.0066 | 0.0001 | -0.05 | 0.12 | 0.65 | <0.0003 |
| #ΔHO | 0% | | | | | |
| †P | 0.99 | | | | | |
| Genes minus tRNA and ribosomal genes | | | | | | |
| All genes | 0.0068 | 0.0001 | -0.20 | 0.08 | 0.66 | <0.0003 |
| CD genes | 0.0069 | 0.0001 | -0.20 | 0.11 | 0.68 | <0.0003 |
| HO genes | 0.0066 | 0.0001 | -0.08 | 0.12 | 0.65 | <0.0003 |
| #ΔHO | -4% | | | | | |
| †P | 0.10 | | | | | |
| Highly expressed genes | | | | | | |
| All genes | 0.0073 | 0.0002 | -0.17 | 0.14 | 0.74 | <0.0003 |
| CD genes | 0.0071 | 0.0002 | -0.07 | 0.17 | 0.75 | <0.0003 |
| HO genes | 0.0077 | 0.0003 | -0.37 | 0.24 | 0.72 | <0.0003 |
| #ΔHO | 8% | | | | | |
| †P | 0.15 | | | | | |
| Highly expressed genes minus tRNA and ribosomal genes | | | | | | |
| All genes | 0.0074 | 0.0002 | -0.30 | 0.17 | 0.73 | <0.0003 |
| CD genes | 0.0072 | 0.0002 | -0.14 | 0.21 | 0.74 | <0.0003 |
| HO genes | 0.0079 | 0.0003 | -0.59 | 0.27 | 0.72 | <0.0003 |
| #ΔHO | 10% | | | | | |
| †P | 0.12 | | | | | |
| Highly expressed genes minus tRNA genes | | | | | | |
| All genes | 0.0074 | 0.0002 | -0.27 | 0.16 | 0.73 | <0.0003 |
| CD genes | 0.0071 | 0.0002 | -0.11 | 0.19 | 0.74 | <0.0003 |
| HO genes | 0.0079 | 0.0003 | -0.59 | 0.27 | 0.72 | <0.0003 |
| #ΔHO | 11% | | | | | |
| †P | 0.08 | | | | | |
| Essential genes | | | | | | |
| All genes | 0.0069 | 0.0002 | -0.38 | 0.28 | 0.70 | <0.0003 |
| CD genes | 0.0067 | 0.0003 | -0.20 | 0.33 | 0.71 | <0.0003 |
| HO genes | 0.0075 | 0.0005 | -0.94 | 0.54 | 0.68 | <0.0003 |
| #ΔHO | 12% | | | | | |
| †P | 0.26 | | | | | |
| ‡Ribosomal genes | | | | | | |
| All genes | -0.00002 | 0.0003 | 2.20 | 0.34 | 0.00008 | 0.96 |
| CD genes | -0.00002 | 0.0003 | 2.31 | 0.36 | 0.0008 | 0.88 |
| HO genes | 0.0073 | 0.015 | -0.66 | 3.44 | 0.11 | 0.82 |
| #ΔHO | NA | | | | | |
| †P | NA | | | | | |

SE = standard error of the estimate of the parameter

$R^2$ = the coefficient of determination, which is the fraction of the variation of the variable, in this case BPSs per CDS, that is explained by the linear model.

*$P_F$ = the probability that the regression occurred by chance, calculated from the F distribution (1) and adjusted for multiple comparisons by the Benjamini–Hochberg procedure (2).

#ΔHO = the % increase of the slope of the HO genes over the slope of the CD genes

†P = the probability that the slopes for CD and HO oriented genes are equal calculated from the two-tailed Student's t distribution (1) and adjusted for multiple comparisons by the Benjamini–Hochberg procedure (2).

‡Included are all genes for ribosomal RNAs and proteins; excluded are all tRNA genes whether or not they are in *rrn* operons. None of the three slopes is different than zero, plus there are only four ribosomal genes in the HO orientation and their CDSs accumulated only 4 BPSs.

CDSs = coding sequences, BPSs = base pair substitutions, indels = insertions and deletions ≤ 4 bp, CD = codirectional with replication, HO = head-on to replication, NA = not applicable.

**Table S4.** Comparisons of the frequencies mutations in the promoters of genes oriented CD versus HO to replication

| | BPSs/promoter | | Indels/promoter | |
|---|---|---|---|---|
| | Mean | SD | Mean | SD |
| All promoters | | | | |
| All genes | 0.14 | 0.42 | 0.04 | 0.41 |
| CD genes | 0.14 | 0.41 | 0.03 | 0.36 |
| HO genes | 0.15 | 0.42 | 0.04 | 0.47 |
| *P | 0.86 | | 0.71 | |
| Promoters of known genes | | | | |
| All genes | 0.14 | 0.40 | 0.04 | 0.39 |
| CD genes | 0.14 | 0.41 | 0.04 | 0.33 |
| HO genes | 0.14 | 0.40 | 0.05 | 0.45 |
| *P | 0.96 | | 0.62 | |
| Promoters of highly expressed Genes | | | | |
| All genes | 0.14 | 0.14 | 0.05 | 0.48 |
| CD genes | 0.14 | 0.14 | 0.03 | 0.31 |
| HO genes | 0.14 | 0.14 | 0.06 | 0.63 |
| *P | 0.92 | | 0.59 | |
| Promoters of essential genes | | | | |
| All genes | 0.13 | 0.40 | 0.05 | 0.47 |
| CD genes | 0.13 | 0.36 | 0.08 | 0.58 |
| HO genes | 0.15 | 0.46 | < 0.01 | NA |
| *P | 0.76 | | NA | |

*P= the probability that the indicated values for CD and HO oriented genes are equal calculated from the Student's two tailed t distribution (1) and adjusted for multiple comparisons by the Benjamini–Hochberg procedure (2). None of the comparisons are statistically significant with or without the adjustment.

BPSs = base pair substitutions, indels = insertions and deletions ≤ 4 bp. CD = codirectional with replication, HO = head-on to replication, SD = Standard deviation; NA = not applicable (no mutations).

**Table S5**. *Bacillus subtilis* mutational data and results

A. Comparison of the mutation results from genes oriented HO versus CD to replication

| | No. CDSs | No. CDSs with BPSs | No. BPSs in CDSs | No. CDSs with indels | No. Indels in CDS | BPSs per CDS | | BPSs/CDS/Nt x $10^3$ | | Indels per CDS | | Indels/CDS/Nt x $10^3$ | |
|---|---|---|---|---|---|---|---|---|---|---|---|---|---|
| | | | | | | Mean | SD | Mean | SD | Mean | SD | Mean | SD |
| All genes | 4,418 | 3,281 | 9,587 | 1,118 | 2,100 | 2.23 | 2.60 | 2.56 | 2.42 | 0.48 | 1.09 | 0.67 | 1.97 |
| CD genes | 3,254 | 2,427 | 7,049 | 786 | 1,476 | 2.17 | 2.61 | 2.45 | 2.42 | 0.45 | 1.08 | 0.57 | 1.65 |
| HO genes | 1,164 | 854 | 2,538 | 332 | 624 | 2.18 | 2.52 | 2.60 | 2.43 | 0.54 | 1.12 | 0.92 | 2.65 |
| *ΔHO | | | | | | 1% | | 6% | | 18% | | 61% | |
| #P | | | | | | 0.92 | | 0.15 | | 0.07 | | <0.0003 | |

B. Correlation of the numbers of BPSs per CDS with the length of the CDS in Nt.

| | Slope | SE | Intercept | SE | $R^2$ | †$P_F$ |
|---|---|---|---|---|---|---|
| All genes | 0.0025 | $3.1 \times 10^{-5}$ | 0.03 | 0.04 | 0.59 | <0.0003 |
| CD genes | 0.0024 | $3.5 \times 10^{-5}$ | 0.03 | 0.04 | 0.59 | <0.0003 |
| HO genes | 0.0027 | $6.7 \times 10^{-5}$ | 0.01 | 0.07 | 0.58 | <0.0003 |
| *ΔHO | 10% | | | | | |
| #P | 0.003 | | | | | |

*ΔHO = the % increase of the values for the HO genes over those of the CD genes

#P= the probability that the indicated values for CD and HO oriented genes are equal from the two-tailed Student's t distribution and adjusted for multiple comparisons by the Benjamini–Hochberg procedure (2). By the nonparametric Mann-Whitney test, the difference between indels in CDSs oriented HO versus CD is statistically significant, P= 0.01 (adjusted).

$R^2$ = the coefficient of determination, which is the fraction of the variation of the variable, in this case BPSs perCDS, that is explained by the linear model.

†$P_F$ = the probability that the regression occurred by chance, calculated from the F distribution and adjusted for multiple comparisons by the Benjamini–Hochberg procedure (2).

### Figure S1

| Promoter | Nt of TSS | Orien-tation | Position relative to TSS | | | | | | | | | | | | | | | | | | | | | | | | | TSS |
|---|---|---|---|---|---|---|---|---|---|---|---|---|---|---|---|---|---|---|---|---|---|---|---|---|---|---|---|---|
| | | | -25 | -24 | -23 | -22 | -21 | -20 | -19 | -18 | -17 | -16 | -15 | -14 | -13 | -12 | -11 | -10 | -9 | -8 | -7 | -6 | -5 | -4 | -3 | -2 | -1 | +1 |
| TSS_671 | 532861 | CD | A | A | C | T | G | G | G | C | T | A | T | A | C | C | G | T | A | G | A | T | A | A | A | G | A | A |
| TSS_1057 | 927728 | HO | A | T | C | G | C | A | C | G | C | A | A | C | T | A | G | A | G | A | A | T | A | C | A | G | A | A |
| asnSp | 988267 | HO | T | G | C | A | G | A | T | G | C | C | A | G | G | T | A | A | C | A | T | A | G | G | T | A | T | C |
| TSS_1387 | 1146994 | CD | C | G | A | C | G | C | C | A | T | C | A | C | G | C | C | A | T | T | A | C | T | T | G | C | T | A |
| ymgFp1 | 1218037 | CD | A | C | A | T | A | C | A | T | C | G | A | C | C | T | G | G | T | C | A | A | A | A | T | T | A | T |
| TSS_1729 | 1341395 | HO | G | A | C | T | T | A | T | T | C | T | T | A | G | C | T | A | T | T | A | T | A | G | T | T | A | T |
| TSS_1874 | 1658536 | HO | A | A | A | A | T | A | T | A | A | C | C | C | A | C | G | A | C | A | A | C | C | A | T | A | A | A |
| TSS_2266 | 2004156 | HO | A | T | C | C | G | A | T | T | A | C | G | G | C | T | A | C | G | C | T | T | C | T | A | A | T | G |
| TSS_2315 | 2077493 | CD | T | C | A | G | C | A | A | G | G | A | T | A | A | A | G | G | G | T | A | T | G | A | T | A | G | T |
| glpABCp | 2350605 | HO | T | T | C | G | A | A | T | T | A | T | G | A | G | C | G | A | A | T | A | T | G | C | G | C | G | A |
| yfaDp5 | 2354712 | HO | T | T | C | G | C | T | G | C | G | A | A | C | A | T | C | C | G | A | T | T | A | C | G | C | T | A |
| dapEp | 2589582 | HO | G | T | G | C | C | C | G | G | T | A | A | G | C | C | T | A | T | G | C | T | G | C | T | G | G | G |
| norRp | 2830329 | CD | T | C | G | A | T | A | T | G | A | C | A | A | T | A | T | C | T | A | T | A | G | T | C | A | A | A |
| ygglp6 | 3087610 | HO | T | G | T | T | T | A | T | C | C | T | G | C | T | G | A | C | A | T | T | G | T | G | G | C | T | G |
| higAp8 | 3232363 | CD | T | G | C | T | A | A | G | G | G | A | T | A | T | T | T | C | A | A | A | A | A | A | C | C | T | G |
| kdsCp2 | 3340032 | HO | T | G | T | C | A | T | T | T | G | C | G | A | T | G | A | C | A | A | T | A | T | G | A | T | G | A |
| rpmBp | 3810017 | CD | C | A | G | T | A | A | A | G | C | G | G | A | T | A | A | A | C | T | C | A | T | T | A | C | T | G |
| rhaDp | 4092407 | HO | T | A | C | A | G | G | T | C | G | G | C | A | A | T | A | G | T | T | G | T | A | G | G | C | C | T |
| acsp1 | 4285617 | HO | G | T | A | A | G | T | G | C | A | T | G | T | A | A | A | A | T | A | C | C | A | C | T | T | T | A |
| pyrLp2 | 4470575 | HO | C | T | T | C | T | G | A | C | G | A | T | G | A | G | T | A | T | A | A | T | G | C | C | G | G | A |
| TSS_5126 | 4518685 | HO | C | G | C | C | A | G | C | T | T | A | G | T | T | T | T | A | T | A | A | T | C | G | G | G | T | T |
| TSS_5192 | 4638448 | HO | G | T | T | G | A | C | G | T | T | G | A | T | G | G | A | A | A | G | T | G | C | A | T | C | A | A |
| Yoshiyama et al 2001 | | | | | | | | | | | | | | | | | | | | | | | | | | | | |
| rpsLp | 3472643 | HO | T | T | T | C | G | G | C | A | T | C | G | C | C | C | T | A | A | A | A | T | T | C | G | G | C | G |

Figure S1. Search for A:T transitions at the 3’ base in -10 elements. Shown are the sequences of promoters in which A:T transitions occurred within 20 Nt of a transcription start site (TSS). Possible -10 elements are boxed, the mutated bases are colored orange, and likely -10 elements with a 3’ T are colored yellow. The mutated -10 element found by Yoshiyama *et al.* (1) is shown at the bottom. The first column gives the promoter designation from Regulon DB (2), the second column gives the position of the TSS (numbered according to NCBI reference sequence NC_000913.2), and the third column indicates whether the promoter is oriented codirectional (CD) or heal-on(HO) to replication.

**Figure S2**

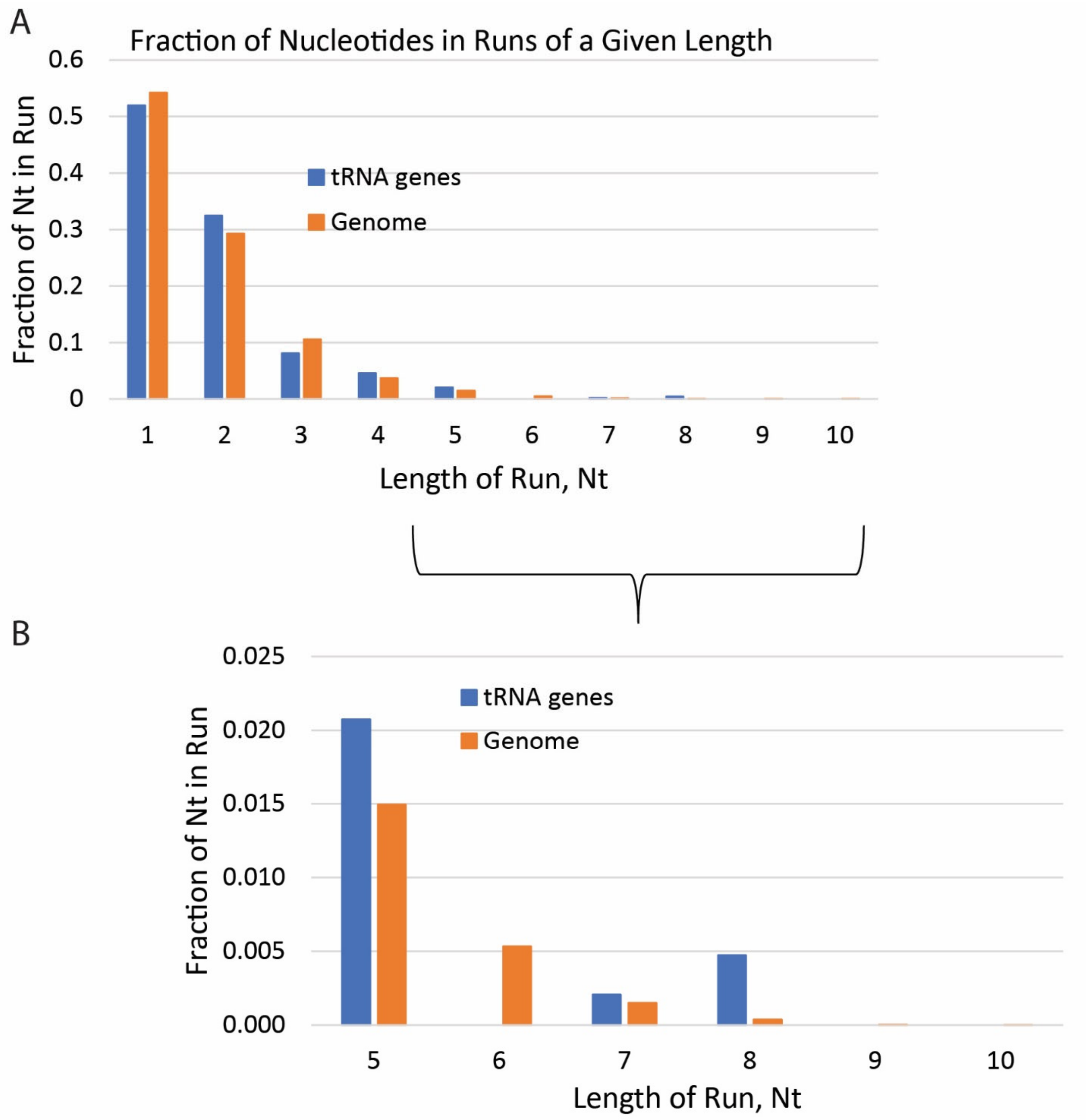


Figure S2. The distribution of mononucleotide runs in tRNA genes and across the genome. A: the number of nucleotides in mononucleotide runs of the length shown was divided by the total number of nucleotides for tRNA genes (blue) and for the entire genome (red). B: expansion of the results for mononucleotide runs of the lengths 5 to 10 Nt showing tRNAs have a higher fraction of runs 5 and 8 Nt long, but not 6, 7, 9, or 10 Nt long.

#### Figure S3

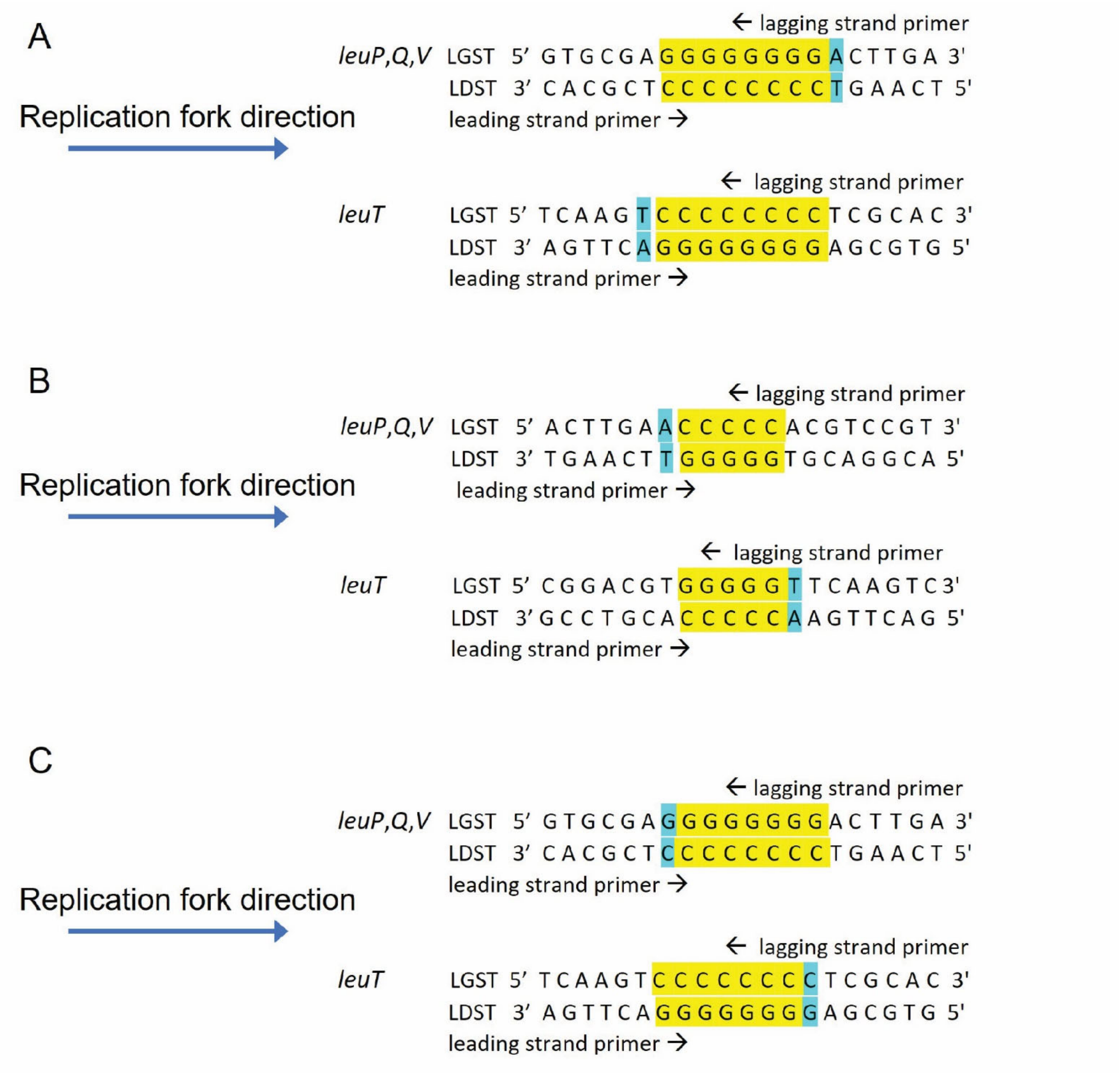


Figure S3. The effects of differences in orientation on possible BPSs due to slipped-mispairing at runs. A: a transition of the A:T bp (indicated in blue) adjacent to the 8 G:C bp run (indicated in yellow) occurred 12 times in *leuP*, *Q*, and *V*, but 0 times in *leuT*. This mutation was likely templated by the run during primer loop-out, which could occur during leading strand replication in *leuP*, *Q*, and *V*, but would have to occur during lagging strand replication in *leuT*. B: a transition of the A:T bp (indicated in blue) adjacent to the 5 G:C bp run (indicated in yellow) occurred once in *leuT* but did not occur in *leuP*, *Q*, or *V*. As in the case illustrated in A, this mutation was likely templated by the run during primer loop-out, which could occur during leading strand replication in *leuT* but would have to occur during lagging strand replication in *leuP*, *Q*, and *V*. C: a transition of the terminal G:C bp (indicated in blue) of the same 8 G:C bp run (indicated in yellow) occurred once in *leuP*. This mutation was likely templated by the adjacent bp during template loop-out, which could occur during lagging-strand replication in *leuP*, *Q*, and *V* but would have to occur during leading-strand replication in *leuT*. LGST = lagging-strand template; LDST = leading strand template.

Figure S4

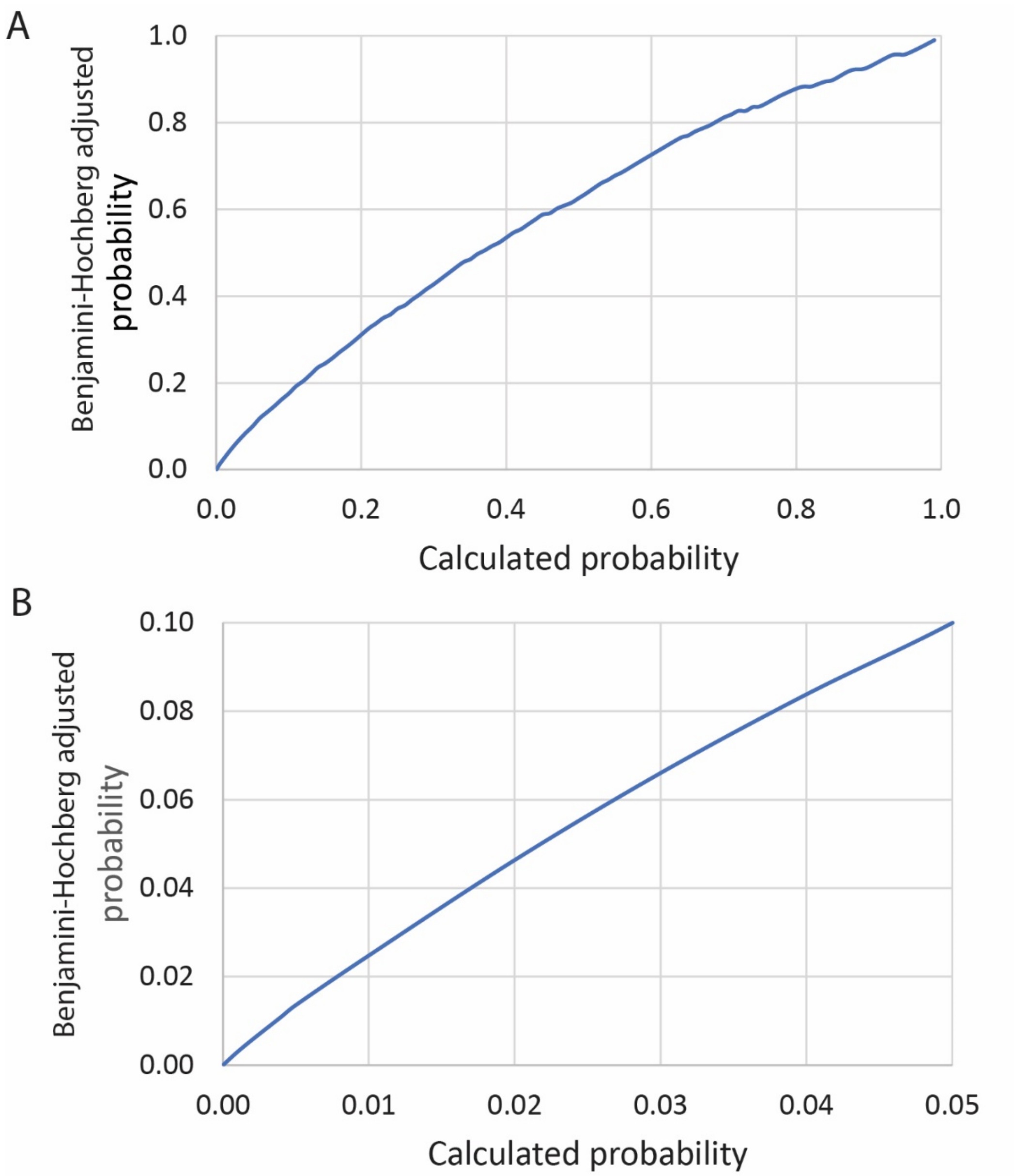


Figure S4. The relationship between the calculated probability values for the statistical tests performed for this paper and the Bonferroni-Holm (1) adjusted probabilities values. A. Calculated probabilities up to a value of 1.00. B. A subset of calculated probabilities up to a value of 0.05. To achieve the accepted significant probability of 0.05 or less after the Bonferroni-Holm adjustment, the calculated probability would have to be ≤ 0.022.
